# Bemnifosbuvir and remdesivir inhibit tick-borne encephalitis virus infection in complementary *in vitro* and *ex vivo* disease models

**DOI:** 10.1101/2025.11.13.688315

**Authors:** Simone Leoni, Isabel Schultz-Pernice, Zuzana Krátka, Luděk Eyer, Lucie Freiholz, Amal Fahmi, Teodora David, Antoinette Golomingi, Fabian Blank, Milan Dejmek, Denis Grandgirard, Ronald Dijkman, Daniel Růžek, Marco P. Alves, Stephen L. Leib

## Abstract

The geographical distribution and incidence of tick-borne encephalitis (TBE) have steeply increased over the past decades, raising to represent a major health concern in Asia and Europe. Symptoms of TBE, caused by infections with tick-borne encephalitis virus (TBEV), range from mild, flu-like symptoms to severe neurological disease, often accompanied by long-term sequelae persisting for several years following pathogen encounter. While effective vaccines against TBEV are available, no antiviral drugs are currently approved and therapeutic options for patients suffering from TBE are limited to supportive measures. Compounds able to disrupt viral nucleic acid synthesis bear the potential of effectively limiting viral replication and spread. Seeking to fill the therapeutic gap, we evaluated the efficacy of a panel of approved and investigational antiviral compounds in containing TBEV infection. Combining several cell lines, human neural organoids and organotypic rat brain slice cultures, we found that the nucleoside analogs remdesivir and bemnifosbuvir efficiently limit viral replication. Through infectious virus quantification, immunofluorescence analysis and flow cytometry, we report significant, dose-dependent reduction of viral loads across all models used, with inhibition observed at low doses for both drugs. Notably, while we observed bemnifosbuvir to be well tolerated, we report important cytotoxicity of remdesivir when applied to human neural organoids. Our findings identify bemnifosbuvir and remdesivir as novel treatment strategies for TBE, providing an accessible and timely response to a clinical challenge of pressing concern.

## INTRODUCTION

Tick-borne encephalitis (TBE) is a severe neurologic disease caused by infections with tick-borne encephalitis virus (TBEV), a member of the *Orthoflavivirus* genus. A steady increase in the incidence of TBE observed across Europe and Asia represents a significant public health concern. TBEV’s increase in geographic range and seasonal incidence are likely related to climate change and land-use alterations expanding tick habitats (*1, 2*). Primarily transmitted through the bite of infected ticks, TBEV can cause a wide spectrum of clinical manifestations, ranging from mild flu-like symptoms to severe neurological complications such as encephalitis, and long-term deficits (*3-5*). Despite the existence of vaccines, treatment for TBE remains supportive, and no specific antiviral therapy is currently approved (*6, 7*). This therapeutic gap contributes to unfavorable outcomes in severe cases and underscores the urgent need for effective treatments. Antiviral strategies that inhibit viral replication early in infection are especially important to prevent central nervous system (CNS) involvement and long-term sequelae. Moreover, the emergence of TBEV genetic variants (*8, 9*) and documented vaccine breakthroughs (*10*) highlight the limitations of current preventive measures and reinforce the necessity for direct-acting antivirals.

Among the most promising candidates for anti-TBEV therapy are nucleoside and nucleotide analogs (NAs), which mimic the natural substrates of viral RNA and DNA synthesis and have proven successful against a range of RNA viruses, such as hepatitis C virus, Ebola virus, SARS-CoV-2, Zika virus, West Nile virus, and dengue virus (*11*). Incorporation of these analogs into the elongating viral RNA molecule results in halting of the viral RNA-dependent RNA polymerase (RdRp), leading to premature chain termination, mutagenesis, or enzyme stalling, thus blocking replication or leading to defective viral proteins, reducing viral load (*12*). To date, only a few NAs have been approved for the treatment of RNA virus-related infections, including sofosbuvir (*13, 14*), ribavirin (*15, 16*), remdesivir (*17-19*), molnupiravir (*20*), and favipiravir (*21*). Some of these compounds have shown successful inhibition of TBEV replication in various cell lines, including A549, Huh7.5, Daoy, and SH-SY5Y (*15, 22, 23*). Additionally, other experimental NAs, such as 3’-deody-3’-fluoroadenosine (*24*), galidesivir (*25*), 2’-C-methyladenosine (*26*), 2’C-methylcytidine (*26*), and 7-deaza-2’-C-methyladenosine (*27, 28*), have demonstrated anti-TBEV activity in preclinical studies.

Driven by the urgent lack of effective TBE therapies, we systematically evaluated a panel of approved and investigational ProTide NAs (*29*) using complementary *in vitro* and *ex vivo* models, including non-neuronal and neuronal cell lines, human neural organoids (hNOs), and organotypic brain slice cultures (OTCs). This strategy enabled the evaluation of antiviral efficacy in physiologically relevant contexts while also revealing tissue-specific tolerability profiles. Importantly, among the tested compounds, we identified remdesivir and bemnifosbuvir as consistently potent inhibitors of TBEV replication, albeit with distinct safety profiles. These findings provide a comprehensive proof-of-concept that repurposed NAs can suppress TBEV replication in pre-clinical models, establishing a foundation for the clinical advancement of direct-acting antivirals against one of Europe’s and Asia’s most problematic neurotropic pathogens.

## RESULTS

### Initial screen identifies several nucleoside analogues inhibiting TBEV replication

To determine which of the selected compounds has an influence on TBEV replication, we tested six different NAs, including sofosbuvir, bemnifosbuvir, uprifosbuvir, and remdesivir (Figure 1). Concurrently to infection, A549 cells, known to efficiently replicate orthoflaviviruses, were treated with each compound at a final concentration of 50 µM, and viral titers were measured by plaque assay two days post-infection (p.i.). Compared with the untreated control, the NAs exhibited differential inhibitory effects on viral replication. Bemnifosbuvir and remdesivir exhibited the strongest antiviral effects, reducing viral titers by approximately 7-log and 6-log respectively. Sofosbuvir led to a 4-log reduction in viral titers, while uprifosbuvir showed a minimal but significant reduction (Figure 1A). In line with these findings, immunofluorescence analysis demonstrated a clear reduction in infected cells after treatment with bemnifosbuvir and remdesivir, while the decrease observed for sofosbuvir was not visually discernible (Figure 1B).

**Figure 1:**
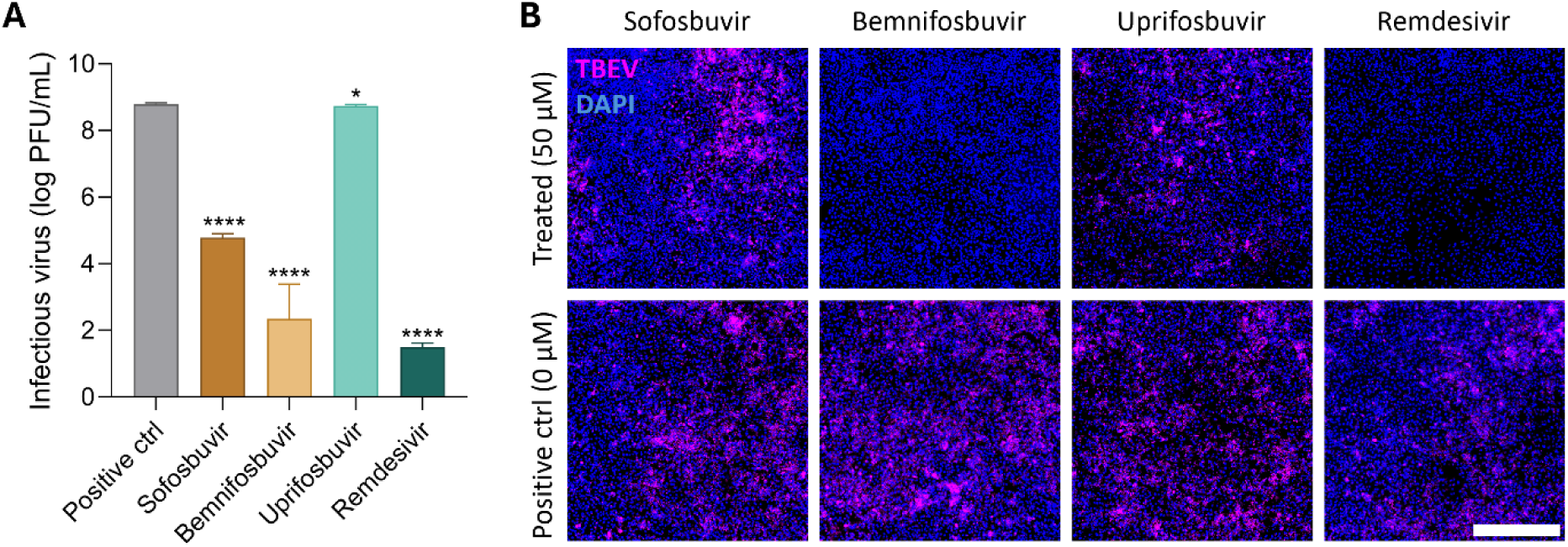
Screening of nucleoside/nucleotide analogs against TBEV in A549 cells. A549 cells were infected with TBEV, treated with selected NAs at 50 µM and incubated for 48 hours. **A,** Virus titers were determined using plaque assays. Data are shown as mean ± s.d. n = 5 independent experiments. Statistical analysis was performed using Welch’s t-test. *p ≤ 0.05, ****p ≤ 0.0001. **B,** A549 cells were fixed on slides, stained with a mouse orthoflavivirus E-protein–specific antibody (magenta), and counterstained with DAPI (blue). Scale bar, 500 μm.

### Nucleoside analogs display dose-dependent activity and cytotoxicity in cell lines

To further investigate the antiviral activity of the selected NAs against TBEV, neuronal (SK-N-SH, SH-SY5Y) and non-neuronal (A549, Vero) cells were concomitantly infected and treated with increasing concentrations (0-50 µM) of sofosbuvir, bemnifosbuvir, uprifosbuvir, and remdesivir. Viral titers were determined two days post-infection, and in parallel, cell viability was assessed to exclude compound-related cytotoxicity. Cell viability assays demonstrated that none of the tested compounds induced significant cytotoxic effects across the concentration range applied. In A549, Vero, and SK-N-SH cells, viability remained stable with increasing compound concentrations (Fig. 2A-C), confirming that the observed reductions in viral titers were attributable to antiviral activity rather than toxicity. A549 cells, bemnifosbuvir and remdesivir exhibited the strongest inhibition of TBEV replication (Fig. 2D), further confirmed by immunofluorescence analysis (Supplementary Fig. 1A). Both compounds showed a clear dose-dependent effect, reducing the viral titers to undetectable levels at concentrations ≥ 6.3 µM. Sofosbuvir displayed only limited activity, with modest reductions in viral replication observed only at the highest concentrations, while uprifosbuvir showed no measurable antiviral effects. The calculated half-maximal inhibitory concentrations (IC_50_) values were 1.39 µM (95% CI: 1.21 – 1.59 µM) for bemnifosbuvir, 1.77 µM (95% CI: 1.55 - 2.02 µM) for remdesivir, and 24.19 µM (95% CI: 20.96-27.91 µM) for sofosbuvir (Fig. 2E; Table 1). In Vero cells, antiviral activity was less pronounced compared to all other cell lines. Remdesivir at high concentrations was the only compound able to reduce viral replication in this cell line (Fig. 2F). The calculated IC_50_ was 14.43 µM (95% CI: 8.80 – 24.24 µM) (Fig. 2G; Table 1). TBEV antigen presence in virus-infected, compound-treated cell culture was confirmed through immunofluorescence microscopy up to the highest concentration of remdesivir tested, confirming only partial replication inhibition (Supplementary Fig. 1B). In neuronal cells, both bemnifosbuvir and remdesivir demonstrated strong antiviral activity, whereas no effect was seen from sofosbuvir and uprifosbuvir at any tested concentration (Fig. 1H, L). In SK-N-SH cells, bemnifosbuvir reduced viral titers below the detection limit at concentrations ≥25 μM, whereas remdesivir required concentrations of at least 50 μM to achieve a comparable effect (Fig. 2H). Immunofluorescence staining of TBEV antigen confirmed this observation (Supplementary Fig. 1C). Undifferentiated SH-SY5Y, a subclone of the SK-N-SH, showed similar values for bemnifosbuvir inhibition with complete inhibition at concentrations ≥ 25 μM (Fig. 1L). A more potent response was observed in differentiated SH-SY5Y (dSH-SY5Y) cells, where bemnifosbuvir suppressed viral replication below the detection limit at concentrations ≥ 12.5 μM, while remdesivir reached this threshold at ≥ 25 μM (Fig. 2L). In SK-N-SH cells, the IC_50_ value of bemnifosbuvir was 4.74 μM (95% CI: 4.10–5.47 μM), compared with 19.50 μM (95% CI: 17.65–21.54 μM; Table 1). In undifferentiated SH-SY5Y, the IC_50_ value for bemnifosbuvir was close to the one of SK-N-SH, with a value of 6.82 μM (95% CI: 6.23 – 7.43 μM) (Fig. 1M; Table 1). In dSH-SY5Y cells, lower IC_50_ values were obtained, indicating a higher sensitivity of these cells to the compounds, with bemnifosbuvir showing an IC_50_ of 1.88 μM (95% CI: 1.62–2.19 μM) and remdesivir of 8.31 μM (95% CI: 7.60–9.08 μM) (Fig. 2M; Table 1). Taken together, these results demonstrate that bemnifosbuvir and remdesivir are the most potent inhibitors of TBEV replication across all tested cell lines, although their efficacy varies depending on the cell type. Bemnifosbuvir was particularly effective in neuronal cells, reducing viral titers below the detection limit at lower concentrations and showing low IC_50_ values. Remdesivir also displayed strong antiviral activity, with consistent inhibition in both neuronal and non-neuronal cells, but at higher effective concentrations compared to bemnifosbuvir. In contrast, sofosbuvir exhibited only modest inhibitory effects in A549 cells, with activity restricted to the highest concentrations tested, while uprifosbuvir did not demonstrate measurable antiviral efficacy in any of the tested cell lines. Importantly, none of the compounds induced significant cytotoxicity under the tested conditions, confirming that the observed viral reduction was specific for the antiviral effect. did not demonstrate measurable antiviral efficacy in any of the tested cell lines.

**Figure 2:**
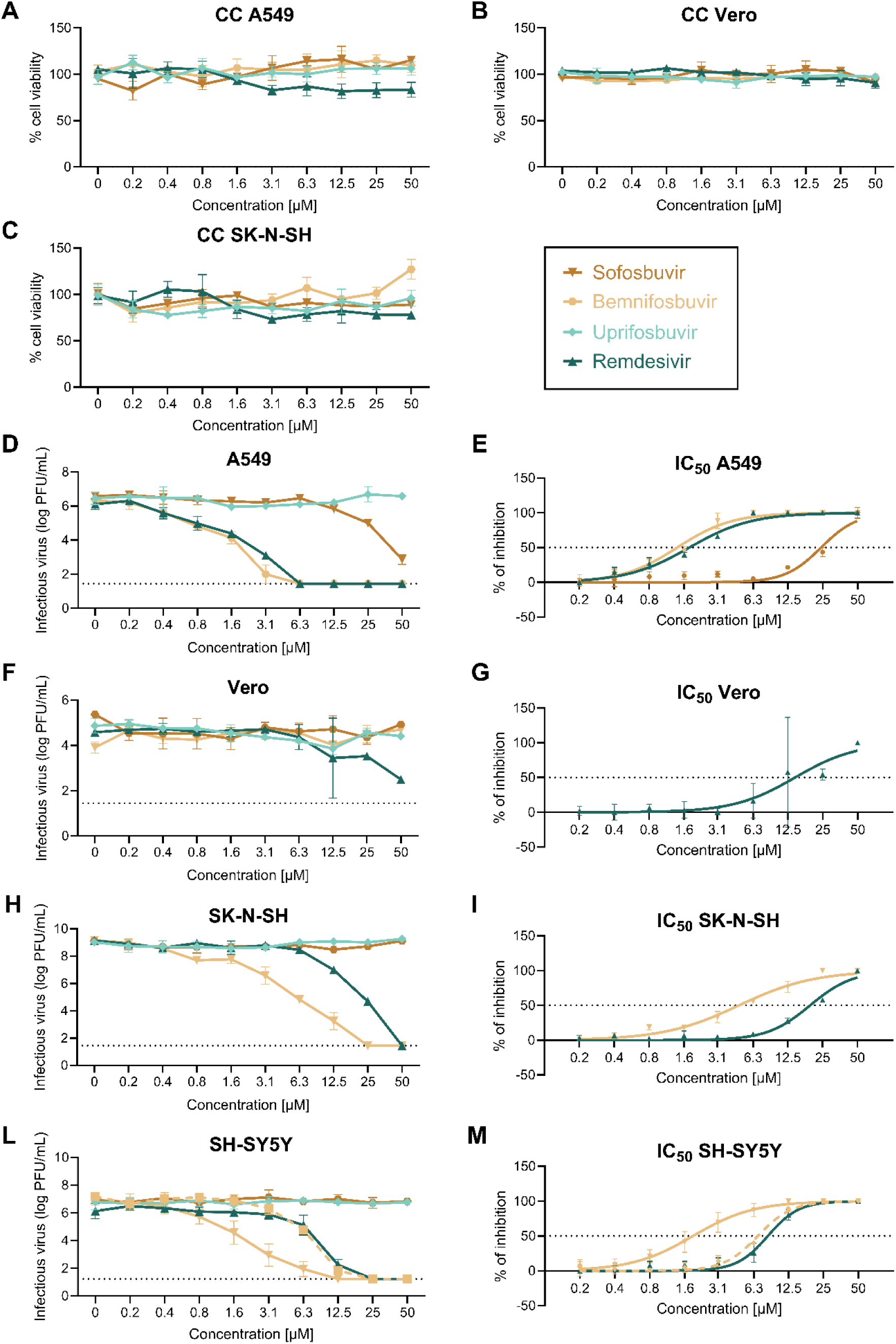
Antiviral activity of nucleoside analogs against TBEV in neuronal and non-neuronal cell lines. **A-C,** Cytotoxicity of the tested compounds measured in terms of cell viability using A549, Vero, and SK-N-SH cells. The cells were treated with increasing concentrations (0-50µM) of uprifosbuvir, remdesivir, sofosbuvir, and bemnifosbuvir and cultivated for 48 hours. **D, F, H, L,** Anti-TBEV activity of the tested compounds was tested by infecting the cells with TBEV. Cells were subsequently treated with increasing concentrations (0-50 µM) and cultivated for 48 hours. Viral titers were determined using plaque assays. Horizontal dotted line indicates the detection limit of the assays. **E, G, I, M,** Inhibition dose-response curves used to calculate the IC_50_ using the nonlinear regression method. The dashed line in panels **L** and **M** indicates values measured in undifferentiated SH-SY5Y cells. Horizontal dotted line indicates 50% inhibition.

**Table 1:**
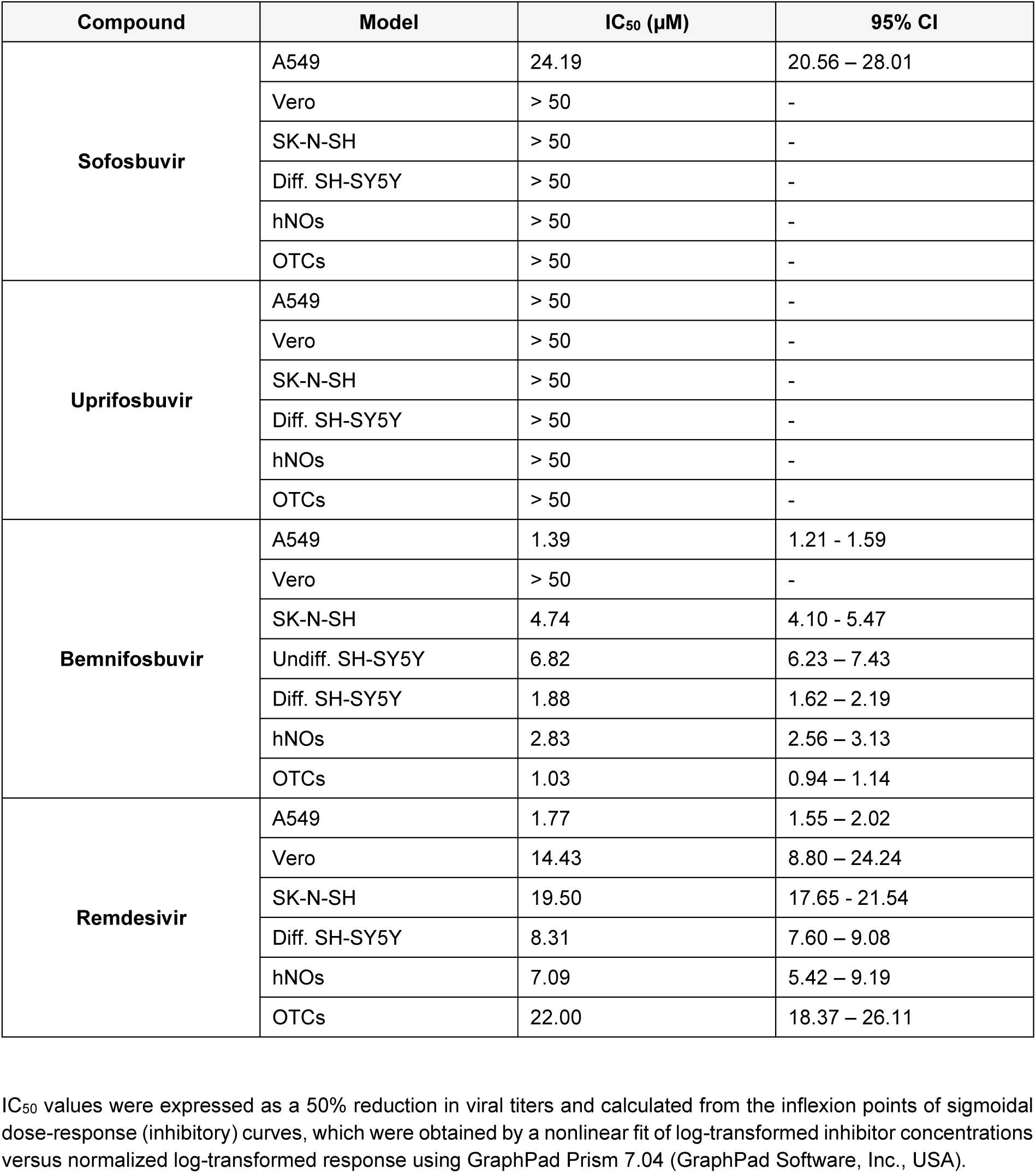
Antiviral and cytotoxicity properties of tested nucleoside prodrugs.

### Remdesivir and bemnifosbuvir reduce infectious TBEV release in human neural organoids

Advanced *in vitro* systems constitute an ethical and scalable platform for drug screening. Aiming to confirm the effect of treatment with the selected NAs in a more complex biological model, we decided to test the activity of the selected antiviral compounds in human neural organoids. To do so, three independent batches of TBEV-challenged organoids were cultured in medium supplemented with 50 μM of uprifosbuvir, remdesivir, sofosbuvir or bemnifosbuvir. Infectious virus released into culture supernatants was quantified at regular intervals for a duration of 8 days and showed important differences between the analyzed NAs (Fig. 3A). While released infectious virus loads peaked at day 2 p.i. in untreated cultures, a strong whilst delayed virus replication was still observed following uprifosbuvir or sofosbuvir treatment, with TBEV titers peaking at day 4 or day 6 p.i. (Fig. 3A). Moreover, while infectious TBEV released from uprifosbuvir-treated cultures followed a shifted but comparable replication curve to the one observed for untreated hNOs, with TBEV titers decreasing between day 4 and day 6 p.i., virus shed from organoids maintained in sofosbuvir-supplemented medium displayed sustained virus release throughout the experiments (Fig. 3A). In contrast, a strong inhibition of infectious TBEV release was documented following remdesivir or bemnifosbuvir treatment, with the latter completely preventing virus replication throughout the whole experiment duration (Fig. 3A). Accordingly, maximum infectious virus titers reached throughout all analyzed time points were significantly reduced in remdesivir and bemnifosbuvir-treated cultures, while no relevant differences were recorded following uprifosbuvir and sofosbuvir supplementation (Fig. 3B). Seeking to analyze possible effects of drug exposure on hNO structure and cells, organoids were processed for immunofluorescence analysis and stained with markers specific to the predominant cell populations, including SOX2-positive neural progenitor cells (*30, 31*) and TUJ1-expressing neurons (*32-34*) (Fig. 3C). Uninfected, mock-treated organoids displayed a lack of envelope (E) protein-harboring cells along with an intact structural and cellular morphology at all analyzed time points (Fig. 3C, Supplementary Fig. 2A and B). In contrast, untreated but TBEV-challenged organoids showed important virus antigen accumulation from day 4 p.i. onwards, along with SOX2 signal waning at day 8 p.i. (Fig. 3C, Supplementary Fig. 2A and B). Similarly, uprifosbuvir and sofosbuvir exposed hNOs displayed strong virus accumulation in neuron-rich organoid areas, while in remdesivir and bemnifosbuvir-treated cultures no viral antigen presence was evident at any analyzed time point (Fig. 3C, Supplementary Fig. 2A and B). Interestingly, sustained SOX2 signal presence was identified in both sofosbuvir and bemnifosbuvir-exposed cultures, while organoids maintained in remdesivir-supplemented medium showed comparable SOX2 signal decline to untreated and uprifosbuvir-treated cultures, indicating heterogeneous effects of NAs addition on neural progenitor cell survival (Fig. 3C, Supplementary Fig. 2A and B).

**Figure 3:**
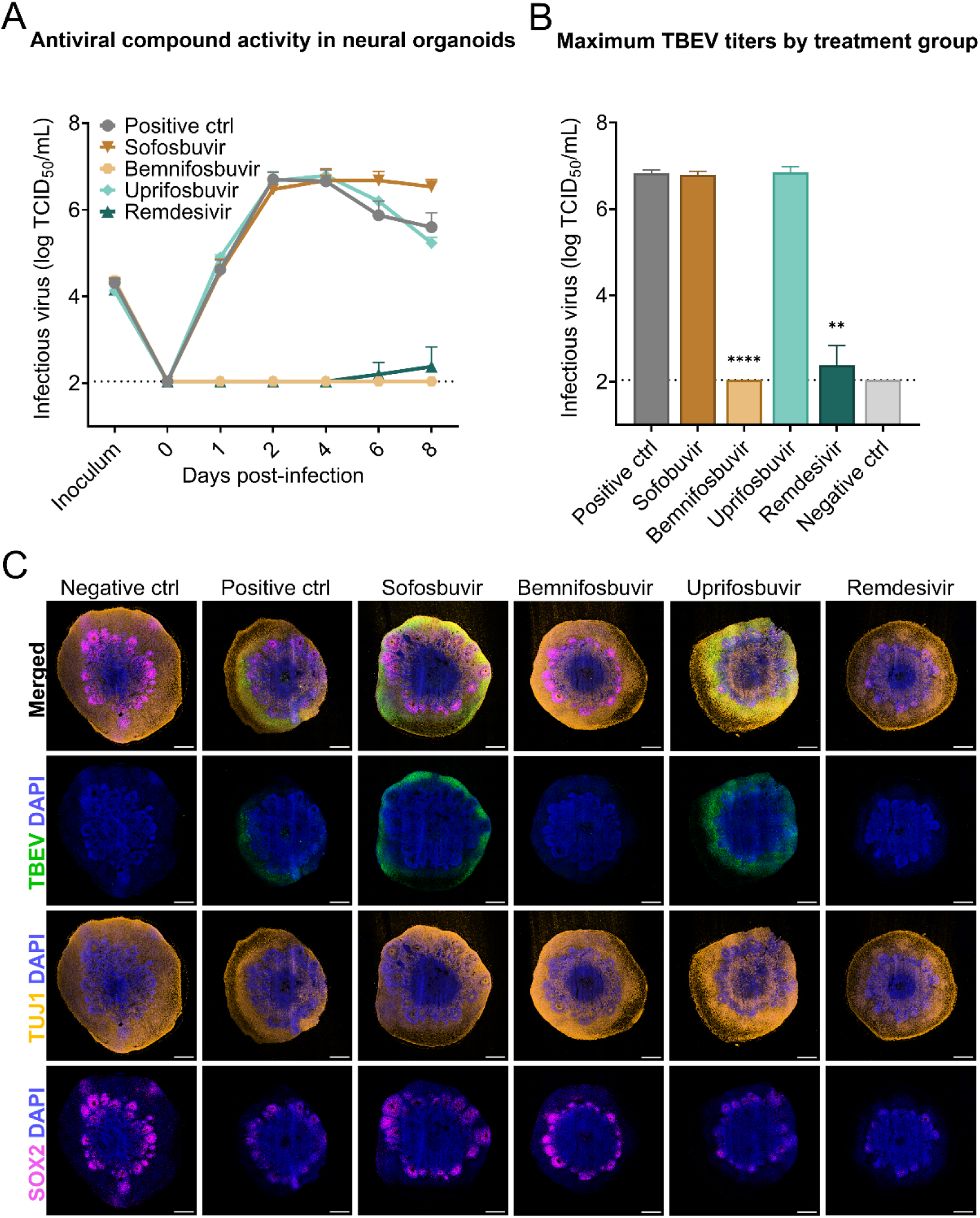
Remdesivir and bemnifosbuvir inhibit TBEV replication in human neural organoids. **A**, Infectious released virus dynamics in TBEV challenged hNOs with or without the addition of NAs at 50 μM. Points indicate mean values ± s.d. n = 3 independent organoid batches. Horizontal dotted line indicates assay detection limit. **B**, Maximum reached TBEV titers across all sampling time points in TBEV-infected hNOs with or without NAs treatment at 50 μM. Bars indicate mean values ± s.d. n = 3 independent organoid batches. Horizontal dotted line indicates assay detection limit. Statistical analysis was performed using Welch’s t-test. **p≤0.01, ****p≤0.0001. **C**, Representative tile-scan confocal micrographs of mock-infected and TBEV-challenged hNOs with or without NA treatment after 8 days of infection. n = 3 independent organoid batches DAPI, blue; TBEV E-protein (4G2), green; TUJ1, orange; SOX2, magenta. Scale bars, 500 µm.

### Remdesivir and bemnifosbuvir show antiviral activity at low doses with variable cytotoxicity in human neural organoids

Seeking to delve into the characterization of dose-dependent TBEV replication inhibition and structural effects exerted by remdesivir and bemnifosbuvir on hNOs, three independent organoid batches were subjected to treatment with the selected NAs at concentrations between 100 μM and 0.39 μM. Bemnifosbuvir-exposed hNOs showed complete suppression of virus replication up to a concentration of 25 μM (Fig. 4A and B). Moderate release of infectious TBEV was observed at a dose of 6.25 μM, while cultures exposed to concentrations of 1.56 μM and 0.39 μM reached maximum titers comparable to untreated, TBEV-challenged controls, despite showing altered release timing (Fig. 4A and B). IC_50_ determination confirmed efficient replication inhibition following bemnifosbuvir treatment, with 50% reduction of infectious viral loads achieved by 2.83 µM (95% CI: 2.56-3.13 µM; Table 1) supplementation (Supplementary Fig. 3). In line with virus quantification findings, immunofluorescence analysis of sectioned hNOs at day 8 p.i. confirmed lack of evident virus antigen signal in cultures treated with up to 1.56 μM of bemnifosbuvir, revealing virus-harboring cells only at concentrations as low as 0.39 μM (Fig. 4C). Intact cell type markers were documented in all hNOs exposed to bemnifosbuvir, independently of detected viral load and drug dose, hinting towards increased cell survival compared to untreated cultures (Fig. 4C). Seeking to further analyze the cytoprotective effect of bemnifosbuvir on hNO-enclosed cells, we performed flow cytometry analysis of dissociated organoids at day 8 p.i., aiming to quantify virus-harboring and apoptotic cells (Fig. 4D and E, Supplementary Fig. 4A). In harmony with our previous findings, flow cytometry quantification of E-protein-positive events showed an evident increase of infected cells in cultures treated with 1.56 μM and 0.39 μM of bemnifosbuvir, while only a small percentage of virus-harboring cells was identified at 6.25 μM (Fig. 4D). The cytoprotective effect of bemnifosbuvir previously suggested by immunofluorescence analysis was confirmed by flow cytometry results (Fig. 4E). Decreased accumulation of cells expressing the apoptotic marker cleaved caspase-3 (CC3) was identified both in mock- and TBEV-challenged cultures in the presence of bemnifosbuvir (Fig. 4E). TBEV-challenged organoids treated with the NA showed a strong drop of apoptotic events, as indicated by a decreased accumulation of CC3-positive cells, reaching percentages comparable to uninfected controls up to a bemnifosbuvir concentration as low as 1.56 μM (Fig. 4E). Moreover, increased cell survival was still detected in organoids cultured in medium supplemented with 0.39 μM of bemnifosbuvir when compared to untreated, TBEV-challenged hNOs, indicating possible beneficial effects of low-dose drug treatment beyond inhibition of virus replication (Fig. 4E).

**Figure 4:**
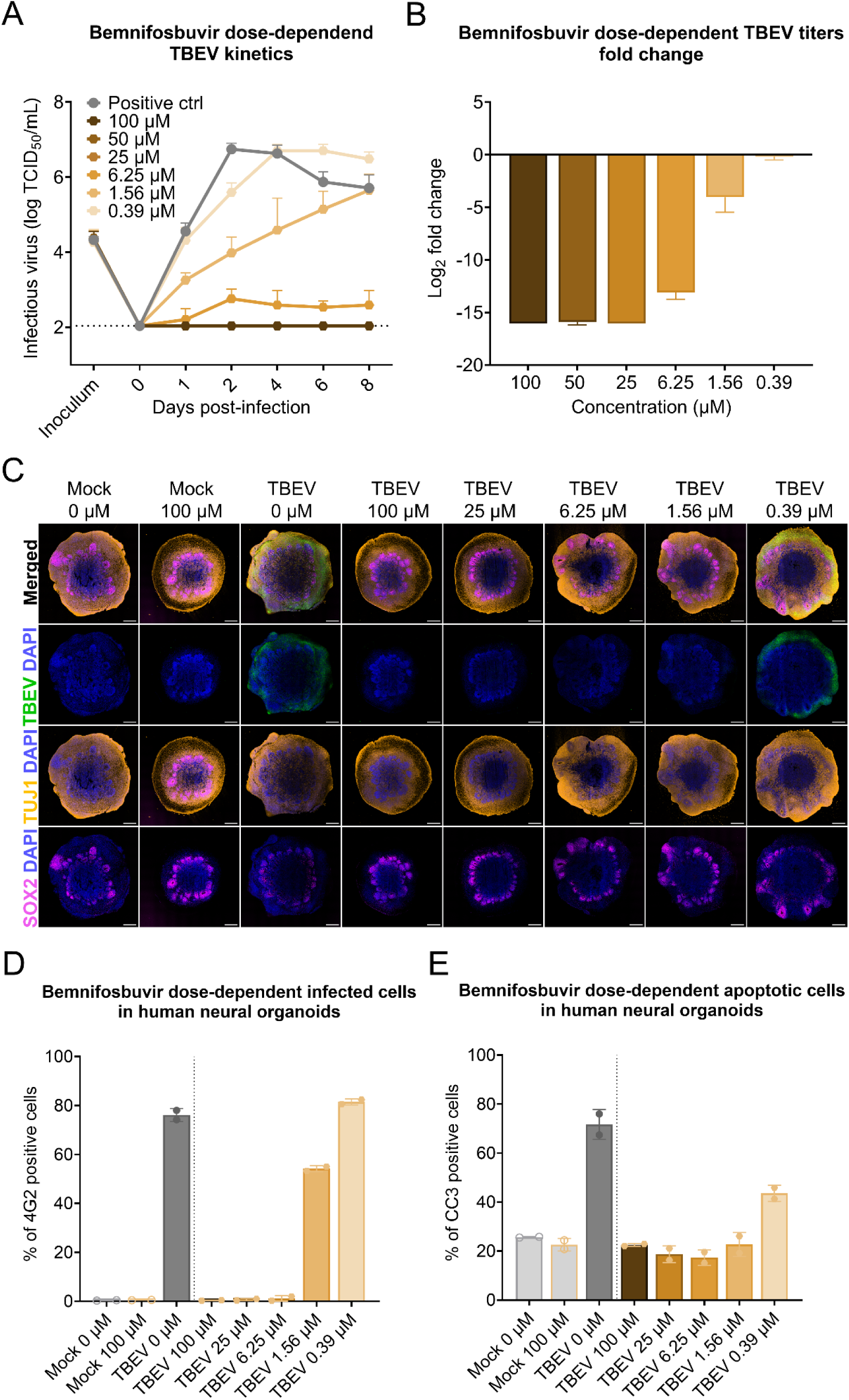
Bemnifosbuvir inhibits viral replication at low doses and displays limited cytotoxicity. **A**, Infectious released virus dynamics in TBEV challenged hNOs with and without addition of 100 μM-0.39 μM bemnifosbuvir. Points indicate mean values ± s.d. n = 3 independent organoid batches. Horizontal dotted line indicates assay detection limit. **B**, Mean Log_2_ fold change of maximum released infectious virus titers following TBEV challenge and treatment with 100 μM-0.39 μM of bemnifosbuvir. Mean values displayed ± s.d. n = 3 independent organoid batches. **C**, Representative tile-scan confocal micrographs of mock-infected and TBEV-challenged hNOs with or without treatment 100 μM-0.39 μM bemnifosbuvir after 8 days of infection. n = 3 independent organoid batches. DAPI, blue; TBEV E-protein (4G2), green; TUJ1, orange; SOX2, magenta. Scale bars, 500 µm. **D**, Percentage of virus-harboring cells detected following 4G2-staining and flow cytometry analysis of TBEV-challenged and mock-infected organoids treated with 100 μM-0.39 μM of bemnifosbuvir after 8 days of infection. Points represent individual organoids (n = 1 independent organoid batch). **E**, Percentage of CC3-positive cells detected through flow cytometry analysis of TBEV-challenged and mock-infected organoids treated with 100 μM-0.39 μM of bemnifosbuvir after 8 days of infection. Points represent individual organoids (n = 1 independent organoid batch).

Inferior potential of remdesivir to contain infectious virus spread compared to bemnifosbuvir was detected through TBEV quantification in remdesivir-treated hNOs (Fig. 5A and B). While concentrations of 100 μM completely prevented infectious virus release, TBEV presence was detected from 50 μM onwards (Fig. 5A and B). Partial replication inhibition was observed in cultures exposed to 50 μM, 25 μM and 6.25 μM of remdesivir, that displayed strongly delayed virus release and dampened titers, while NA concentrations of 1.56 μM and 0.39 μM showed limited effect in restraining TBEV spread (Fig. 5A and B). Accordingly, IC_50_ assessment confirmed 50% virus replication inhibition at 7.09 µM (95% CI: 5.42 – 9.19 µM) (Table 1) remdesivir treatment (Supplementary Fig. 3). Immunofluorescence microscopy of hNOs sectioned after 8 days of infection, revealed clear virus accumulation starting at concentrations of 6.25 μM of remdesivir, with high amounts of virus-harboring cells evident at 1.56 μM and 0.39 μM (Fig. 5C). Furthermore, cell marker analysis confirmed the previously observed fading of SOX2 signal following remdesivir exposure independently of virus challenge (Fig. 5C). Near to complete ablation of neural progenitor cell marker fluorescence was observed in hNOs exposed to 100 μM, 50 μM and 25 μM of remdesivir, while SOX2 signal was restored in organoids treated with 6.25 μM and 1.56 μM of the drug (Fig. 3C and 5C). Flow cytometry analysis of hNOs at day 8 p.i. further substantiated observed virus dynamics and cell death induction (Fig. 5D and E). Increasing percentages of virus antigen-harboring cells were detected through flow cytometry starting at remdesivir concentrations of 25 μM, with E-protein-positive cells reaching comparable levels to untreated cultures when exposed to 1.56 μM and 0.39 μM of the NA (Fig. 5D). In line with previous findings, strong apoptosis induction was confirmed following hNO remdesivir exposure up to a drug concentration of 25 μM, independently of mock- or TBEV-infection (Fig. 5E). In particular, uninfected hNOs maintained in medium supplemented with 100 μM of remdesivir displayed mean apoptotic cell percentages of 59.6%, compared to 25.7% detected in untreated mock organoids (Fig. 5E). Similarly, following TBEV-challenge and concomitant remdesivir treatment, cultures exposed to 100 μM of the NA comprised 53.1% apoptotic cells, while 35.8% of cells were found to express CC3 in organoids subjected to 25 μM drug treatment (Fig. 5E). To confirm distribution of apoptotic cells and assess whether SOX2 signal disappearance following remdesivir treatment correlated with neural progenitor cell death, organoids collected 8 days p.i. were sectioned and analyzed through fluorescence microscopy following CC3-staining (Supplementary Fig. 4B and C). Immunofluorescence imaging confirmed limited apoptosis in bemnifosbuvir-exposed cultures (Supplementary Fig. 4B), while signal increase was evident in remdesivir-treated hNOs independently of virus exposure (Supplementary Fig. 4C). Moreover, in untreated TBEV-challenged organoids and both uninfected and infected organoids exposed to 100 μM and 25 μM of remdesivir, CC3 signal was observed accumulating in organoid regions displaying ventricular morphology, confirming increased, selective death of neural progenitor cells (Supplementary Fig. 4C). Taken together, our results indicate that, while both bemnifosbuvir and remdesivir can potentially limit infectious TBEV spread, possible cell toxicity need to be taken into consideration when treating patient populations.

**Figure 5:**
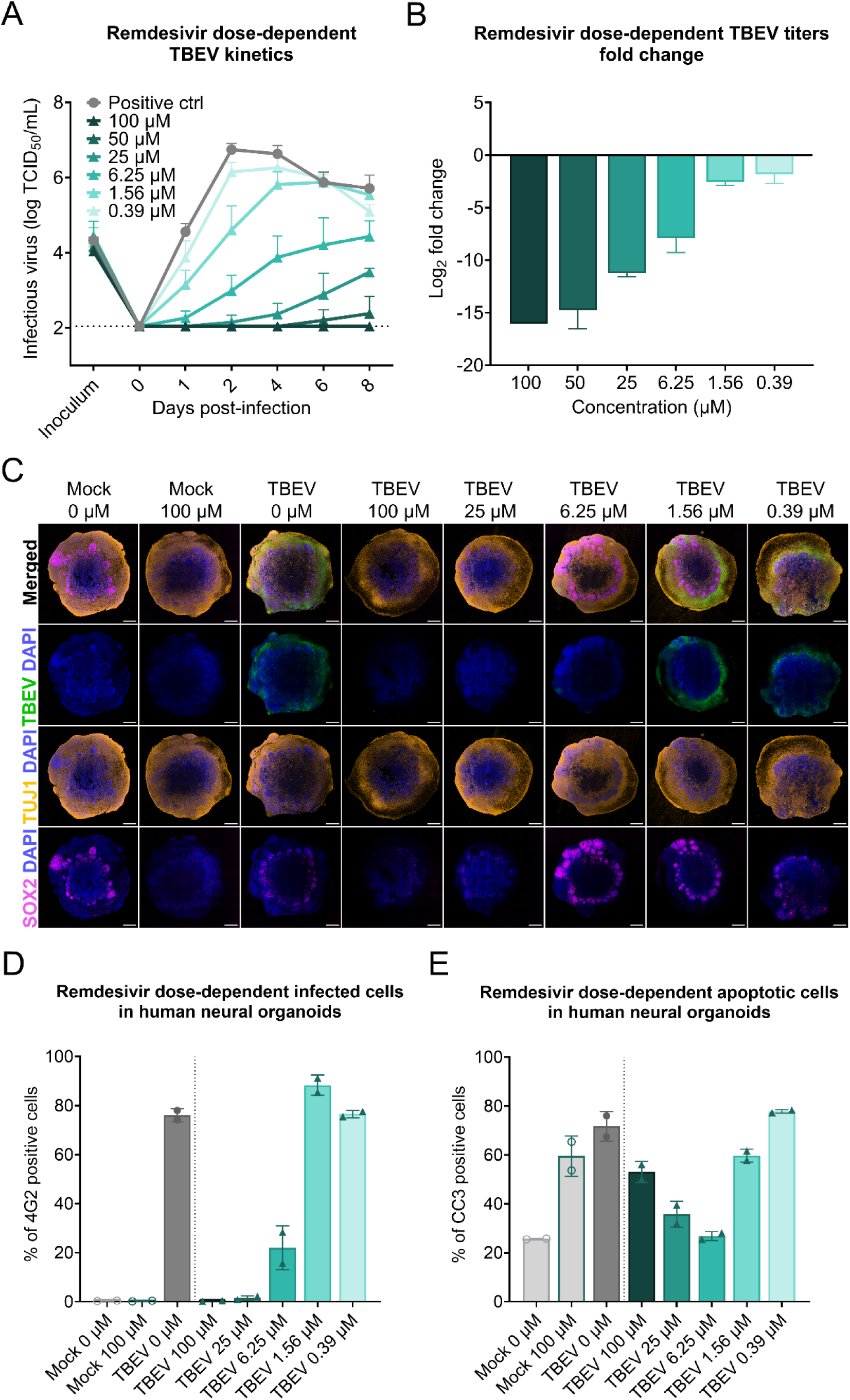
Remdesivir inhibits viral replication in a dose-dependent manner but leads to increased neural progenitor cell apoptosis. **A**, Infectious released virus dynamics in TBEV-infected hNOs with and without the addition of 100 μM-0.39 μM remdesivir. Points indicate mean values ± s.d. n = 3 independent organoid batches. Horizontal dotted line indicates assay detection limit. **B**, Mean Log_2_ fold change of maximum released infectious TBEV titers following hNO infection and treatment with 100 μM-0.39 μM of remdesivir. Mean values displayed ± s.d. n = 3 independent organoid batches. **C**, Representative tile-scan confocal micrographs of mock-infected and TBEV-challenged organoids with or without treatment 100 μM-0.39 μM remdesivir at 8 days p.i. n = 3 independent organoid batches. DAPI, blue; TBEV E-protein (4G2), green; TUJ1, orange; SOX2, magenta. Scale bars, 500 µm. **D**, Percentage of virus-harboring cells detected by 4G2-staining and flow cytometry analysis of TBEV- and mock-infected hNOs treated with 100 μM-0.39 μM of remdesivir at 8 days p.i. Points represent individual organoids (n = 1 independent organoid batch). **E**, Percentage of CC3-positive cells detected through flow cytometry analysis of TBEV- and mock-infected hNOs treated with 100 μM-0.39 μM of remdesivir at 8 days p.i. Points represent individual organoids (n = 1 independent organoid batch).

### Remdesivir and bemnifosbuvir reduce infectious TBEV release in rat cerebellar slice cultures

To better assess the translational potential of candidate antivirals in system more closely resembling the *in vivo* setting, we evaluated the efficacy of selected NAs in rat cerebellar OTCs from cerebellum exposed to TBEV. OTCs were infected and concomitantly treated and infectious viral titers were assessed three days post infection with a TCID_50_ assay. In a preliminary screening of all antivirals at a concentration of 50 µM, treatment with bemnifosbuvir and remdesivir resulted in significant reduction in viral titers compared to infected DMSO-treated controls. Notably, bemnifosbuvir reduced titers below the detection limit (4-log reduction), indicating a more potent antiviral activity in this *ex vivo* model compared to remdesivir (3-log reduction). In contrast, sofosbuvir and uprifosbuvir failed to suppress viral replication at this concentration. Uprifosbuvir-treated OTCs infected with TBEV did not differ from the DMSO-treated control, while treatment with sofosbuvir showed significantly higher titers, indicating a possible enhancement of viral replication (Fig. 6A). These results were further substantiated by immunostaining for viral antigen and co-staining with Purkinje cells, a known target of TBEV infection (*35, 36*). TBEV-infected cultures treated with uprifosbuvir and sofosbuvir did not differ from the infected control condition, while remdesivir and bemnifosbuvir showed complete depletion of virus antigen signal, comparable to the uninfected control (Fig. 6B).

**Figure 6:**
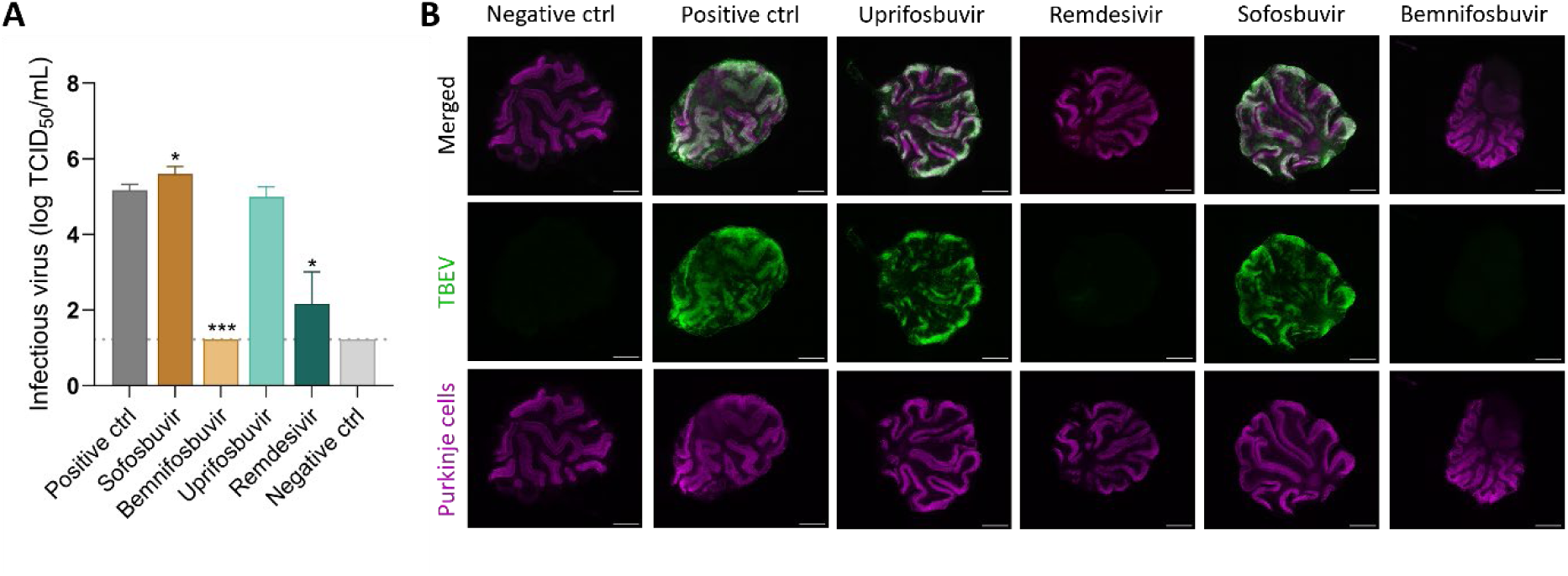
Screening of nucleoside analogs against TBEV in rat brain slice cultures of cerebellum. **A**, Cerebellar brain slice cultures were infected with TBEV and simultaneously treated with selected NAs at 50 µM. Data are shown as mean ± s.d. n = 3 independent experiments. Statistical analysis was performed using Welch’s t-test. *p<0.05, ***p<0.001. **B**, Immunofluorescence staining of cerebellar brain slice cultures at 3 days p.i. TBEV E-protein (T036), green; Purkinje cells/Calbindin, purple. Scale bars, 1 mm.

### Remdesivir and bemnifosbuvir suppress TBEV replication in rat cerebellar slice cultures at low concentrations

To adhere to the 3Rs principle and reduce animal use, follow-up dose-response testing in OTCs was limited to the most effective compounds, i.e. bemnifosbuvir and remdesivir. Both compounds reduced TBEV replication in a concentration-dependent manner, but with marked differences in potency. Bemnifosbuvir achieved complete suppression of viral replication at concentrations as low as 5 µM, whereas remdesivir required 100 µM to reduce viral titers below the detection limit (Fig. 7A). Consistently, the IC_50_ for bemnifosbuvir was 1.03 µM (95% CI: 0.94 – 1.14 µM), while remdesivir showed a substantially higher IC_50_ of 22.00 µM (95% CI: 18.37 – 26.11 µM) (Fig. 7B) (Table 1). Immunofluorescence analysis confirmed the quantitative results at the tissue level. In untreated, infected slices, viral antigen was abundantly distributed throughout the cerebellar cortex, with prominent infection of Purkinje cells. At low antiviral concentrations, staining patterns were comparable to the control condition. Treatment with 5 μM bemnifosbuvir was sufficient to eliminate detectable TBEV antigen (Fig. 7C), whereas for remdesivir, complete clearance required 100 μM; at 50 μM, viral antigen was reduced but still visible (Fig. 7D). These observations were consistent with the viral titration data, further highlighting the greater potency of bemnifosbuvir in this *ex vivo* model. In summary, bemnifosbuvir showed markedly greater potency than remdesivir in OTCs, suppressing TBEV replication and eliminating viral antigen at substantially lower concentrations.

**Figure 7:**
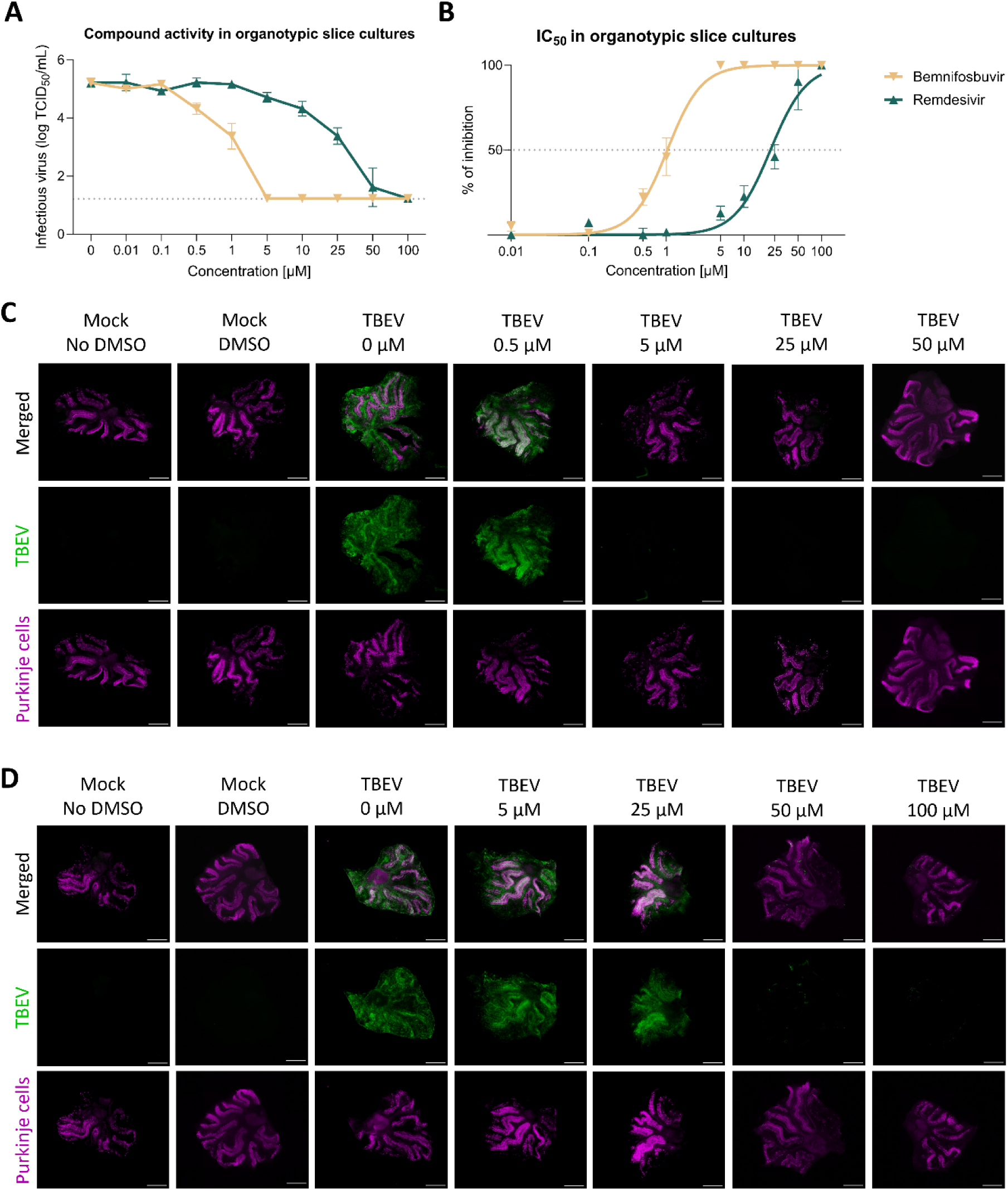
Antiviral activity of remdesivir and bemnifosbuvir in rat cerebellar slice cultures. **A**, Viral titers determined 3 days p.i. showing the inhibition of TBEV replication in OTCs treated with remdesivir and bemnifosbuvir (0–50 µM). Horizontal dotted line indicates the detection limit of the assays. **B**, Dose-response curves used to calculate the IC_50_ using nonlinear regression method. Horizontal dotted line indicates 50% inhibition. **C-D**, bemnifosbuvir treatment in C and remdesivir treatment in D. Immunofluorescence staining of cerebellar brain slice cultures at 3 days p.i. TBEV E-protein (T036), green; Purkinje cells/Calbindin, purple. Scale bars, 1 mm.

## DISCUSSION

Despite the availability of effective vaccines, the growing incidence of TBE and the absence of antiviral therapies demand urgent identification of therapeutics able to mitigate disease progression. NAs have recently emerged as promising candidates with the potential of limiting *Orthoflavivirus*-mediated infections both *in vitro* and *in vivo* (*37-40*). In this study, we employed a range of complementary *in vitro* and *ex vivo* approaches to evaluate the antiviral activity of repurposed NAs, namely sofosbuvir, bemnifosbuvir, uprifosbuvir, and remdesivir against TBEV. All four compounds feature the McGuigan phosphoramidate moiety at the 5′ position, classifying them as members of the McGuigan ProTide family (*41*). Sofosbuvir and bemnifosbuvir share key structural features, most notably the 2′-α-fluoro-2′-β-methyl substitution on the ribose, differing only in their nucleobase: sofosbuvir incorporates uracil, whereas bemnifosbuvir contains guanine (*42, 43*). Uprifosbuvir is structurally similar to sofosbuvir, with the 2′-α-chloro-2′-β-methyl modification distinguishing it (*44*). In contrast, remdesivir is an adenosine-based C-nucleoside, featuring a 1′-cyano group, a C-glycosidic bond, and 4-aza-7,9-dideaza modification of the adenine moiety (*45, 46*). We show that among the tested compounds, bemnifosbuvir and remdesivir consistently demonstrate the strongest inhibitory activity across several cell lines, hNOs, and rat organotypic OTCs. We find that bemnifosbuvir bears superior efficacy in inhibiting TBEV replication, achieving low micromolar IC_50_ values in several systems and reducing viral titers below detection limits at low concentrations. In contrast, we report that remdesivir, while showing broad effectiveness in containing virus spread, displays generally higher IC_50_ values in all neural models, possibly reflecting differences in compound uptake, activation by metabolization of the prodrug or polymerase affinity. We furthermore reveal differences in the tolerability of bemnifosbuvir and remdesivir in hNOs, the latter displaying increased apoptosis induction of neural progenitor cells when applied at high concentrations.

In an initial screen using A549 cells, a commonly used model for orthoflavivirus studies, we identified remdesivir, bemnifosbuvir, and sofosbuvir as the most potent compounds limiting TBEV replication. Unexpectedly, uprifosbuvir was observed enhancing TBEV replication, and was thereby included in further analysis. The antiviral efficacy at several concentrations of these compounds was subsequently evaluated in additional cell lines of both neuronal and non-neuronal origin. In A549 cells, both bemnifosbuvir and remdesivir exhibited potent, dose-dependent inhibition. Sofosbuvir displayed measurable activity only at high concentrations, while uprifosbuvir showed no significant effect on viral replication. In contrast, Vero cells were broadly refractory to the tested compounds. Only remdesivir displayed slight inhibition at high concentrations, while bemnifosbuvir, sofosbuvir, and uprifosbuvir failed to show any effect. In neuronal cell lines (SK-N-SH and SH-SY5Y), a more relevant model of TBEV’s target cells, bemnifosbuvir and remdesivir emerged as the most active compounds in limiting TBEV replication, while no effect was observed following uprifosbuvir and sofosbuvir treatment.

Our study indicates that, although sofosbuvir, bemnifosbuvir, and uprifosbuvir share a high degree of structural similarity, they differ markedly in antiviral activity, likely due to several factors. Previous reports suggest that the active triphosphate form of bemnifosbuvir is generally a more favorable substrate for orthoflaviviral RdRp than sofosbuvir (*47*), probably because its larger guanine base enhances binding to the polymerase active site due to base-stacking interactions, promoting efficient incorporation and subsequent chain termination. Accordingly, sofosbuvir exhibits relatively low antiviral activity against most vector-borne orthoflaviviruses, in line with the findings of this study. In addition, bemnifosbuvir, as a guanine derivative, has been reported to potently inhibit orthoflaviviral NS5 methyltransferase (MTase) (*47, 48*), which likely further contributes to its superior antiviral profile compared to sofosbuvir. Structurally related uprifosbuvir, however, shows little to no antiviral effect against TBEV, probably due to the 2′-α-chloro modification. By contrast, remdesivir’s antiviral potency is conferred by its C1′-cyano substitution (*46*), distinguishing it mechanistically from the 2′-modified ProTides.

Furthermore, the documented differences in antiviral activity could also be explained by cell-dependent features of the prodrugs’ metabolism, as some cell lines have been observed displaying increased efficiency at converting the same prodrug than others (*49*). Prodrug activation occurs in a two-step process. The first step is the transformation of the prodrug to its intermediate metabolite (monophosphate). This reaction is catalyzed by carboxylesterase 1 (CES1), cathepsin A (CatA), and histidine triad nucleotide-binding protein 1 (HINT1) (*49, 50*). The second step is the conversion of the monophosphate to the pharmacologically active triphosphate, which requires sequential phosphorylation by host kinases (*51*). The efficiency of this activation pathway varies among cell types and is superior particularly in primary hepatocytes and hepatocyte-derived cell lines, such as HepG2 and Huh-7 (*52*). In line with our findings, it was shown that CES1 is not expressed in Vero E6 cells (*53*), providing a potential explanation for the reduced effect of the prodrugs. In contrast, A549 cells were reported to show high expression levels of key metabolic enzymes, as well as the relevant drug transporters and organic anion-transporting polypeptides (*53*). Neuronal cell lines may also express elevated levels of these activating enzymes, supporting more efficient prodrug conversion and consequently higher antiviral potency, which is consistent with the strong efficacy observed for bemnifosbuvir and remdesivir in SK-N-SH, SH-SY5Y, and other tested neural models. However, the expression levels of these key enzymes have not yet been extensively studied in neuronal cell types.

The activation of ProTides is further complicated by the pronounced stereospecificity of key enzymes such as CES1 and CatA, which efficiently hydrolyze only one of the two possible diastereomers (*50, 52*). McGuigan ProTides possess a chiral center at the 5′-phosphate, existing as Sp or Rp diastereomers or as a racemic mixture (*41, 50, 52*). Although this stereochemical dependence is largely irrelevant *in vivo*, where multiple cell types contribute to ProTide activation, it can result in false-negative outcomes during antiviral screening if an inappropriate cell line is employed, since the efficacy of activation strongly depends on the expression profile of the relevant enzymes in that particular line. While cell lines represent a scalable and cost-effective approach for high-throughput drug screens, their simplicity allows limited insights into disease and treatment dynamics in the human host. Aiming to further explore the potential of preselected NAs in inhibiting TBEV replication in a more complex system, drug treatment was applied to hNOs, an advanced *in vitro* model of human neural tissue. In line with results obtained in the neuronal cell lines, bemnifosbuvir and remdesivir showed superior efficacy in limiting virus replication compared to uprifosbuvir and sofosbuvir, which yielded no relevant change in viral load. Subsequent dose-defining experiments confirmed higher potency of bemnifosbuvir compared to remdesivir, as previously documented in neuronal cells. Complete inhibition of infectious TBEV release was documented for bemnifosbuvir-treated hNOs at concentrations up to and including 25 µM. In contrast, remdesivir-supplemented cultures allowed virus replication at late time points following exposure to 50 µM of the drug. Cell diversity encompassed in hNOs additionally revealed selective apoptosis induction in neural progenitor cells following remdesivir treatment, demonstrated by dose-dependent CC3 upregulation localized to ventricular areas, concomitant with SOX2 signal loss. NA-induced toxicity has been largely reported both *in vitro* and *in vivo* and has been shown to greatly vary depending on the permeability and metabolism of the exposed cell types, tested compounds and treatment duration (*54, 55*). While remdesivir has been reported to display limited off-target toxicity in several human cell lines (*55*), interactions with mitochondrial DNA- and RNA-polymerases leading to altered mitochondrial gene expression and functions have been suggested as possible mechanisms underlying cytotoxicity following drug exposure, alongside general cell toxicity (*55-61*). Mitochondrial dynamics and metabolism have been shown to play crucial roles in orchestrating neural progenitor cell proliferation and neurogenesis (*62-64*). Moreover, mitochondrial dysfunction during neurogenesis has been linked to loss of neural progenitor cell self-renewal capacity with consequent cell pool depletion and cognitive decline (*65*). Disruption of those tightly controlled mechanisms by treatment of neural progenitor cells with high concentrations of remdesivir, could provide a potential explanation for the observed deleterious consequences of drug treatment in hNOs. Moreover, orthoflaviviruses have been reported to alter mitochondrial morphology and functions, including homeostasis, immune regulation, and apoptosis (*66*), potentially resulting in synergistic effects at intermediate remdesivir concentrations. While remdesivir has been shown not to cross the blood-placental barrier in rats, its main metabolite GS-441524 can be transferred transplacentally (*67*). Despite cytotoxicity being observed at concentrations exceeding remdesivir’s IC_50_ value and above the reported C_max_ of 9 µM (*68*), our findings warrant further investigations on the safety profile of remdesivir in certain vulnerable populations, including pregnant women and neonates.

To further evaluate the four most promising compounds in a highly complex and physiologically relevant setting, we employed organotypic rat OTCs from cerebellum. This *ex vivo* model preserves the cytoarchitecture and cell–cell interactions of the CNS and has previously been used for antiviral screening against neurotropic viruses (*28*). Furthermore, it allows testing of several conditions using the brain of a single animal, reducing the number of animals required for the experiments, in agreement with the 3Rs principle. In accordance with the results obtained in cell lines and hNOs, an initial screen at 50 µM demonstrated measurable inhibition of TBEV replication only upon remdesivir and bemnifosbuvir treatment, whereas sofosbuvir and uprifosbuvir showed no efficacy. Subsequent dose-response experiments were therefore performed only with remdesivir and bemnifosbuvir. In this system, bemnifosbuvir consistently displayed higher potency than remdesivir, achieving stronger suppression of viral replication across a broader range of concentrations. Structural analysis of OTCs processed for immunofluorescence microscopy did not reveal any evident disruption or morphological change of the Purkinje cell layer induced by high-concentration treatment of antivirals. From a translational perspective, the *ex vivo* results are particularly noteworthy. They demonstrate that bemnifosbuvir can achieve complete viral clearance in partially intact neuronal tissue, where drug diffusion barriers, cell-cell interactions, and neuronal susceptibility differ markedly from monolayer cultures. This suggests that the compounds retain efficacy in experimental systems that approximate the biological environment of the infected brain. The superior potency of bemnifosbuvir over remdesivir in OTCs may reflect differences in metabolic activation, polymerase affinity, or intracellular stability of the triphosphate form in a complex system characterized by the interplay of diverse cell populations, as discussed above.

Effective TBE treatment relies on antivirals capable of inhibiting viral replication while reaching therapeutic concentrations in the CNS. In this regard, the observation that TBEV can enter the brain without an early breakdown of the blood-brain barrier (BBB) represents a core challenge. In a murine model, BBB permeability increased only at later disease stages (*69*), implying that barrier disruption is not a prerequisite for TBEV neuroinvasion. This was further confirmed by human BBB models, which show that endothelial cells support TBEV infection and permit transcellular passage of the virus without overtly compromising tight-junction integrity, reinforcing the notion that CNS entry can precede BBB failure (*70, 71*). Therefore, therapeutics may need to cross an initially intact BBB to be effective early in the disease. From this perspective, bemnifosbuvir and remdesivir represent two promising candidates. Their pharmacokinetic properties, however, highlight different translational challenges.

Bemnifosbuvir demonstrated good tolerability in phase I studies (*72, 73*). Although its CNS penetration has not yet been assessed, its oral administration and sustained plasma levels raise the possibility of achieving therapeutic concentrations in the brain, considering that low concentrations of this compound sufficed to inhibit TBEV replication in our models. Remdesivir must be administered intravenously to achieve sufficient concentrations in systemic circulation and reach target cells; when given orally, it is trapped in the liver to be rapidly converted in hepatocytes to its active triphosphate form (*74-76*) (Li et al., 2021, Schäfer et al., 2022, Hu et al., 2020). However, available data suggest that remdesivir achieves only low, although sustained, concentrations in the brain compared to other analyzed tissues (*45*).

Another important consideration is the biphasic clinical course of TBE. After an initial incubation period, patients often present with a first phase characterized by nonspecific, flu-like symptoms, followed by a short remission before the onset of the second, neurological phase (*4*). Antivirals are likely to be most effective if administered during the early viremic phase, before the virus has entered the CNS. However, in clinical practice, patients presenting with flu-like symptoms are typically not tested for TBEV unless there is a specific indication, such as a history of tick bites or a relevant epidemiological context. It is only when patients seek medical care following the onset of neurological symptoms, at which point the virus has already entered the CNS, that specific diagnostic testing is performed. This represents a major challenge for the development of antiviral therapy against TBEV and emphasizes the importance of identifying compounds with sufficient CNS penetration, able to be effective during the second phase of the disease, when viral replication persists within the brain parenchyma (*35, 77*). Since most current nucleotide ProTides, including bemnifosbuvir and remdesivir, were primarily developed to treat viral infections with the predominant site of replication in hepatocytes (e.g., HCV, Ebola virus) next-generation therapeutics should employ novel ProTide strategies that enhance resistance to rapid hepatic metabolism, thereby ensuring high systemic stability, efficient penetration of the BBB, and selective targeting of neuronal cell types. Evidence of viral persistence in CNS during the symptomatic phase underscores the need for therapies that can reduce viral load even after neuroinvasion has occurred. Future studies, aiming at identifying optimal therapy onset timing and compound efficacy at inhibiting viral replication at later infection time points will shed light on possible therapeutic approaches.

Overall, our study highlights the potential of multisystem drug screening approaches in analyzing the efficacy and limitations of prospective therapeutic compounds. By combining a series of cell lines, complex *in vitro* and physiological *ex vivo* models, we highlight the importance of applying relevant experimental systems, as shown by the diverse results obtained in cell line screens. Furthermore, our study emphasized the potential of harnessing integrated model strategies to mitigate technical variables while shedding light on unique aspects based on cell composition, structural and physiological variables of each system. While our multi-model approach strengthens the robustness of our findings, several limitations must be acknowledged. Despite the ability of hNOs and OTCs to recapitulate key aspects of brain development, architecture and cellular diversity, they lack a complete immune compartment, limiting their capacity to integrate other relevant pathological processes. Moreover, neither model reproduces systemic pharmacokinetics, metabolism, and BBB dynamics. Our study did not include *in vivo* experiments, which remain essential to define therapeutic efficacy, drug distribution, and safety in the context of an intact immune system. Finally, although remdesivir and bemnifosbuvir emerged as the most promising candidates, key aspects such as CNS penetration and optimal dosing regimens remain unresolved and warrant further preclinical and clinical evaluation.

In conclusion, our findings demonstrate that NAs can effectively suppress TBEV replication in diverse *in vitro* and *ex vivo* models, including neuronal cell lines, hNOs and OTCs. Across all three models, bemnifosbuvir consistently emerges as the most potent and best-tolerated compound, by demonstrating robust antiviral activity in both hNOs and rat OTCs. In comparison, our study unveils that remdesivir also exhibits potent efficacy, albeit with reduced tolerability. The distinct toxicity profile of remdesivir compared to bemnifosbuvir highlights the importance of compound selection when targeting neural tissue. Overall, this study provides strong pre-clinical evidence that direct-acting antivirals can be used against TBEV. These results constitute an initial step towards potential therapeutic options for TBE. As TBE incidence continues to rise and patients face lifelong neurological sequelae, our work underscores the urgent need to translate these findings into *in vivo* validation and clinical development.

## MATERIAL AND METHODS

### Cell culture

Vero cells (ATCC CCL-81; African green monkey kidney epithelial cells), SK-N-SH cells (ATCC HTB-11; human neuroblastoma cell line), and A549 cells (ATCC CRM-CCL-185; human alveolar basal epithelial cells derived from a male patient with non-small cell lung carcinoma) were cultured in DMEM, supplemented with 10% newborn calf serum or FBS, with or without addition of 100 U/mL penicillin, 100 µg/mL streptomycin, and 1% L-glutamine (all from Sigma-Aldrich). Cells were maintained at 37°C in a humidified atmosphere containing 5% CO_2_. SH-SY5Y (ATCC CRL-2266; Thrice cloned subline of the neuroblastoma cell line SK-N-SH) were cultured in high glucose DMEM containing GlutaMAX (Invitrogen), 10% heat-inactivated FBS, 1 mM sodium pyruvate (Invitrogen), and 1x penicillin-streptomycin (Invitrogen). The embryonic stem cell line H1 (WiCell, WA01, kindly provided by Dr. Marisa Jaconi, University of Geneva, Switzerland) was used for hNO generation. Cells were grown in feeder-free conditions in Vitronectin XF (Stem Cell Technologies) coated flasks with mTeSR Plus medium (Stem Cell Technologies) as per manufacturer’s instructions. Cells were passaged on a weekly basis using a 0.5 mM EDTA (Thermo Fisher Scientific) in Dulbecco’s phosphate-buffered saline (DPBS) solution to obtain small cell aggregates. Mycoplasma absence was confirmed through routine testing using the LookOut Mycoplasma qPCR Detection Kit (Merck). All media and reagents were allowed to reach room temperature prior to usage.

### SH-SY5Y neuronal differentiation

Differentiation was performed in 3 steps: maintenance medium was exchanged for glutamine-free MEM with non-essential amino acids (Invitrogen) supplemented with 5% heat-inactivated FBS, 1x penicillin-streptomycin, 2 mM GlutaMAX, and 10 µM all-trans retinoic acid (Sigma Aldrich) and cells incubated for one week. Then, the concentration of heat-inactivated FBS was reduced to 2% for 3 further days. Finally, cells were seeded on poly-D-Lysin coated vessels in terminal differentiation medium consisting of glutamine-free neurobasal medium (Invitrogen) supplemented with 1x B27 supplement (Thermofisher), 20 mM potassium chloride (Thermofisher), 1x penicillin-streptomycin, 2 mM GlutaMAX, 50 ng/mL brain-derived neurotrophic factor (Sigma Aldrich), 2 mM dibutyryl cyclic AMP (StemCell Technologies), and 10 µM all-trans retinoic acid. Cells were incubated for 6 days before infections were performed.

### Virus propagation

The prototypical TBEV strain “Hypr” (kindly provided by Spiez Laboratory, Swiss Federal Office for Civil Protection, Spiez, Switzerland and by the collection of Arboviruses at the Institute of Parasitology, Biology Centre of the Czech Academy of Sciences, České Budějovice, Czech Republic), belonging to the Western European subtype, was used for all experiments. Upon receival, the virus was passaged up to four times on Vero, A549 or SK-N-SH cells to produce stocks necessary for the present study. Briefly, on the day preceding infection, cells were seeded in cell culture flasks at a density of 6-7 x 10^4^ cells/cm^2^ in Dulbecco’s Modified Eagle Medium (DMEM), supplemented with 10% fetal bovine serum (FBS, Sigma Aldrich or Gibco) and incubated overnight at 37°C with 5% CO_2_. On the following day, medium was replaced with DMEM supplemented with 2% FBS and the required amount of virus suspension was added directly into the flasks. Supernatant was harvested on the third day of infection, aliquoted and stored at -70°C or -80°C. Mock stocks were simultaneously produced, adhering to the same method with omission of virus inoculation. Experiments were performed in biosafety level 2 (Czech Republic) or 3 (Switzerland) conditions in full compliance with local biosafety regulations, by trained staff equipped with appropriate personal protective equipment. Prior to usage, all solutions were allowed to reach room temperature.

### Compound reconstitution

Sofosbuvir and remdesivir were purchased from Biosynth, AA block and Selleck Chemicals. Bemnifosbuvir was purchased from Medchemexpress, Advanced ChemBlocks or Selleck Chemicals and uprifosbuvir from TargetMol. All compounds were solubilized in 100% dimethyl sulfoxide (DMSO) to yield 10 mM stock solutions.

### Cytotoxicity assay

To evaluate the potential cytotoxic effects of the tested compounds, Vero, SK-N-SH, or A549 cells were seeded into 96-well plates at a density of 2×10^4^ cells per well. After a 24-hour incubation period, allowing the formation of a confluent monolayer, the culture medium was carefully aspirated and replaced with fresh DMEM supplemented with the compounds at a range of concentrations (0 - 50 µM). Cells were then incubated at 37°C for an additional 48 hours. Cell viability was assessed using the Cell Counting Kit-8 (Dojindo Molecular Technologies) following the manufacturer’s instructions. The assay provided a quantitative measure of compound-induced cytotoxicity based on metabolic activity.

### Viral titer reduction assay

Vero, A549, and SK-N-SH cells were seeded into 96-well plates at a density of 2×10^4^ cells per well and incubated for 24 hours to allow cell attachment and monolayer formation. For initial screening of antiviral activity, the confluent cell monolayers were infected with TBEV at a multiplicity of infection (MOI) of 0.1 PFU/cell. Immediately following infection, the cells were treated either with a fixed concentration of the test compounds (50 µM) or with a range of concentrations (0-50 µM), and incubated at 37°C for an additional 48 hours post-infection (p.i.).

Viral titers were quantified using plaque assay methodology. The resulting titration data were used to plot dose-response curves, from which the IC_50_ values were calculated based on percent inhibition of viral replication. SH-SY5Y cells (undifferentiated or differentiated according to section "SH-SY5Y neuronal differentiation") were seeded into 48-well plates (2×10^5^ cells/well). Cell monolayers were infected with an inoculum containing both the antiviral compounds at different concentrations range (0-50 µM) and TBEV at a MOI of 0.01 TCID_50_/cell for 30 minutes at 37°C and 5% CO_2_. The inoculum was then substituted with fresh medium supplemented with the respective antiviral following incubation for 48 hours. Infectious viral titers were quantified using a TCID_50_ assay. The obtained data were used to plot dose-response curves, from which the IC_50_ were calculated.

### Plaque assay

Plaque assays were performed using A549 cells, following a modified version of a previously described protocol (*26*). Briefly, serial 10-fold dilutions of the virus were prepared and added to 24-well tissue culture plates, and A549 cells were subsequently seeded at a density of 0.6-1.5×10^5^ cells per well. After a 4-hour incubation at 37°C to allow for virus absorption, an overlay was added, consisting of 1.5 % (w/v) carboxymethyl cellulose in DMEM. The plates were incubated for 5 days at 37°C, after which the monolayers were gently washed with phosphate-buffered saline (PBS) and stained with naphthalene blue-black to visualize plaques. Plaques were subsequently counted, and viral titers were calculated and expressed as plaque-forming units per milliliter (PFU/mL).

### Tissue culture infectious dose assay

One day prior to titration, Vero cells were seeded on 96-well plates at a density of 2 x 10^4^ cells/well in the previously described medium (see section “Cell culture maintenance”) and incubated overnight at 37°C with 5% CO_2_. On the following day, confluency was confirmed, and culture medium was replaced with DMEM supplemented with 2 or 10% FBS. Culture supernatant samples were added to row A to a final dilution of 1:10 and the medium was thoroughly mixed by repeated pipetting. Serial 10-fold dilutions were subsequently obtained by repeated vertical volume transfer. The infection was allowed to proceed for 3 days at 37°C with 5% CO_2_. On day 3, medium was removed by aspiration and plates were fixed with 4% PFA for 20 minutes at room temperature. Plates were subsequently washed three times with 0.1% saponin or buffer composed of sodium chloride, potassium chloride, monopotassium phosphate, di-sodium hydrogen phosphate dihydrate and Tween 20 dissolved in dH_2_O (henceforth cumulatively referred to as permeabilization buffer) and subsequently incubated for up to 45 minutes at 37°C with a monoclonal pan-flavivirus anti-E protein antibody obtained from hybridoma cells (4G2, clone D1-4G2-4-15, ATCC, HB-112), diluted 1:10 in a 0.3% saponin in PBS solution. Following incubation, plates were washed twice with permeabilization buffer, then incubated at 37°C with a 1:250 horseradish-peroxidase conjugated secondary antibody (DAKO #P0260) in 0.3% saponin solution. After a maximum of 45 minutes of incubation, plates were washed twice with permeabilization buffer and cells stained by addition of 3-amino-9-ethylcarbazole (AEC) substrate solution, composed of 50 mM acetate buffer (pH = 0.5), AEC concentrate (1 tablet of 3-amino-9-ethylcarbazol dissolved in 2.5 mL of *N,N*-Dimethylformamide) and 30% hydrogen peroxide. The colorimetric reaction was allowed to proceed for up to 45 minutes, after which the plates were washed with tap water. Viral titers were determined following the Reed and Muench calculation method (*78*) and expressed as TCID_50_/mL.

### Immunofluorescence staining in cell lines

Vero, SK-N-SH, or A549 cells were seeded into 96-well plates at a density of 2 × 10⁴ cells per well, infected with TBEV at a MOI of 0.1 PFU/cell, and treated with test compounds at concentrations of up to 50 μM. The cells were then incubated at 37 °C for 48 hours. Following incubation, cells were fixed and permeabilized using ice-cold acetone–methanol solution (1:1) and subsequently blocked with 10% FBS to reduce nonspecific binding. Immunostaining was performed using a mouse pan-flavivirus anti-E protein antibody (clone D1-4G2-4-15; Sigma-Aldrich; MAB10216-I; 1:250), followed by incubation with an AlexaFluor 647-conjugated goat anti-mouse secondary antibody (A-21235, Invitrogen; 1:1000). Cell nuclei were counterstained with 4′,6-diamidino-2-phenylindole (DAPI, 1 μg/mL). Fluorescent signal detection and image analysis were carried out using the ImageXpress Pico automated imaging system in combination with CellReporterXpress software (Molecular Devices).

### Generation of human neuronal organoids

Human neural organoids were produced as previously published (*79*). For embryoid body generation, on day 0, H1 embryonic stem cells were dissociated into a single cell solution using Accutase (Stem Cell Technologies) and counted. An appropriate volume of cell suspension, equivalent to 10’000 cells/well, was pelleted at 200 g for 5 minutes at 4°C. The supernatant was removed, cells resuspended in the required volume of Formation Medium (STEMdiff Cerebral Organoid Kit, Stem Cell Technologies) supplemented with 50 µM Y-27632 (Stem Cell Technologies) and distributed in ultra-low attachment 96-well plates (Corning), 100 µL per well. Cultures were incubated at 37°C, with 5% CO_2_ and 100 µL of Formation Medium was added in two-day intervals. Dissociated embryonic stem cells were furthermore used for stemness assessment via flow cytometry analysis. To this purpose, a volume equivalent to ∼1 x 10^6^ cells was centrifuged at 280 g for 8 minutes at 4°C. Supernatant was aspirated and cells resuspended in 100 µL of antibody solution composed of SSEA-5 (Stem Cell Technologiesand TRA-1-60 (Stem Cell Technologies) antibodies suspended in BD CellWASH (BD Biosciences). Following a 15-minute incubation period in the dark, on ice, cells were washed with 1 mL of BD CellWASH and spun down at 380 g for 8 minutes at 4°C. The supernatant was removed, and the pellet resuspended in 100 µL of BD CellWASH. Samples were acquired using a FACS Canto II (BD Bioscience) in combination with the DIVA software. Flow cytometry data was analyzed using FlowJo 10.9.0 (ThreeStar). A stemness of minimum 80% was deemed sufficient to proceed with organoid generation. On day 5, successful embryoid body formation and culture quality was visually assessed, and the formed structures transferred to ultra-low attachment 24-well plates (Corning) containing 500 µL of pre-warmed Induction Medium each (STEMdiff Cerebral Organoid Kit, Stem Cell Technologies). Cultures were incubated two more days at 37°C, with 5%CO_2_. On day 7, embryoid body quality was visually assessed and aggregates were individually embedded into 30 µL of ice-cold Matrigel (Stem Cell Technologies). Organoid-harboring Matrigel droplets were transferred into ultra-low attachment 6-well plates (Corning), prepared with 3 mL of pre-warmed home-made Expansion Medium each. For Expansion Medium preparation, DMEM/F-12 (Gibco) and Neurobasal Medium (Gibco) were mixed at a 1:1 ratio and supplemented with 1:100 B27 minus vitamin A (Gibco), 1:100 GlutaMAX (Gibco), 1:200 MEM-NEAA (Seraglob), 2.5 µg/mL of insulin (Sigma-Aldrich), 3.5 µL/L 2-mercaptoethanol (Sigma-Aldrich) and 1:100 Penicillin-Streptomycin. Following mixture, medium was sterile filtered. On day 10 of culture, organoids were visually assessed for sufficient bud formation and structural integrity, and medium was replaced with 3.5 mL of home-made Maturation Medium, containing a 1:1 mixture of DMEM/F-12 (Gibco) and Neurobasal Medium (Gibco) supplemented with 1:100 B27 (Gibco), 1:100 GlutaMAX (Gibco), 1:200 MEM-NEAA (Seraglob), 2.5 µg/mL of insulin (Sigma-Aldrich), 3.5 µL/L 2-mercaptoethanol (Sigma-Aldrich), 1:100 Penicillin-Streptomycin and 0.5 µg/mL Amphotericin B (Gibco). Maturation Medium was sterile filtered prior to usage and kept for a maximum of 14 days. From day 10 on, plates were placed on an orbital shaker moving at a speed of 65 rpm and maintained at 37°C, with 5% CO_2_. Regular medium changes were performed in 2- to 3-day intervals. All reagents and media were allowed to reach room temperature prior to application to the cultures.

### Infection of human neuronal organoids

Organoids of 42 days of development were used for TBEV challenge experiments. One day prior to infection, 4 representative organoids were dissociated in Accutase, and cell numbers were quantified using CountBright Absolute Counting Beads, for flow cytometry (Thermo Fisher) as per manufacturer’s protocol. Samples were analyzed using a FACS Canto II with DIVA software and quantifications were made using FlowJo 10.9.0. For hNO infection, a MOI of 0.01 TCID_50_/cell was used, calculated based on the previously estimated cell numbers. The required volume of Maturation Medium (see “Generation of human neural organoids”) was supplemented with the calculated amount of TBEV Hypr stock and an appropriate volume of NA solution (prepared as per “Compound reconstitution”) or DMSO (equivalent to the highest DMSO concentration reached in NA-treatment conditions). The solution was thoroughly mixed, culture supernatant was removed, and 3 mL of virus or mock suspension were added to each well. Mock infections were carried out analogously, substituting the virus stock with an equivalent volume of uninfected cell supernatant. The infection was allowed to proceed for 2 hours at 37°C, 5% CO_2_ shaking at 65 rpm. Subsequently, the inoculum was removed through aspiration and hNOs were washed three times with 3 mL of DPBS to remove excess virus. Following the last wash, 4 mL of fresh Maturation Medium supplemented with an appropriate amount of NA solution or DMSO, was added to each well and cultures were placed back at 37°C, 5% CO_2_ shaking at 65 rpm. Supernatant and tissue samples were collected at pre-defined intervals and complete medium changes were performed every two days, replacing the culture medium with Maturation Medium freshly supplemented with an appropriate volume of NA-solution or DMSO. All media and reagents were allowed to reach room temperature prior to usage.

### Cryopreservation and immunofluorescence staining of human neural organoids

For organoid cryopreservation and immunofluorescence staining, the previously published protocol was applied (*80*). Briefly, hNOs were washed twice with DPBS and subsequently fixed 10 minutes with 2% PFA in PBS solution followed by a minimum 24-hour fixation in 4% PFA (Formafix). After fixation, organoids were washed twice with DPBS and transferred to a 30% sucrose (Sigma-Aldrich) in DPBS solution. Cultures were left to incubate in the solution at 4°C until hNOs were observed sinking to the container’s bottom, confirming successful equilibration. Organoids were subsequently transferred to a pre-warmed gelatin solution, composed of 7.5% gelatin and 10% sucrose in DMEM, and incubated for a minimum of 1 hour at 37°C, to allow gelatin penetration. For snap-freezing, hNOs were transferred into Cryomolds (Biosystems Switzerland AG) with an appropriate volume of gelatin solution and gently placed to float on a mixture composed of 100% ethanol on dry ice. Frozen hNOs were stored at -80°C until further processing. For immunofluorescence staining, organoid-harboring gelatine blocks were processed into 18 µm thin slices using a Leica CM1950 cryostat and collected on SuperFrost Plus Adhesion slides (Epredia). Slides were stored at -20°C until staining. For immunofluorescence staining, selected slides were allowed to reach room temperature. To ensure gelatin removal, slides were subsequently incubated 10 minutes in a 0.1% Tween-20 in PBS solution (PBS-T) pre-heated to 37°C. Staining areas were encircled using a ReadyProbes Hyrophobic Barrier Pap Pen (Thermo Fisher Scientific) and sections blocked with a 5% goat serum in PBS-T for 1 hour at room temperature in a moisturized chamber.

Primary antibodies (Table 1A) were diluted in a 5% bovine serum albumin in PBS-T solution and subsequently applied onto the hNO sections following blocking completion. Primary antibody incubation was allowed to proceed overnight at 4°C in a moisturized chamber. On the following day, slides were washed three times in PBS-T for 5 minutes to ensure unbound antibody removal. Secondary antibodies (Table 1B) were diluted in PBS-T and the solution was distributed on the organoid sections for 2 hours at room temperature in a moisturized chamber. Following secondary antibody incubation, slides were washed three times for 5 minutes in PBS-T and mounted using MS Shield Mount with DABCO (Electron Microscopy Sciences). Confocal imaging was performed at the microscopy imaging center (MIC) of the University of Bern, Switzerland, using a Carl Zeiss LSM710 microscope. All micrographs were adjusted equally between experiments for optimal brightness and contrast using imageJ 2.14.0/1.54f.

### Flow cytometry analysis of human neural organoids

For flow cytometry analysis, hNOs were dissociated by 10-minute incubation in 500 µL of Accutase at 37°C followed by manual dissociation through repeated pipetting. 250 µL of the obtained suspension, estimated to contain approximately 1.5 x 10^6^ cells, were transferred to a new tube and the Accutase was diluted by the addition of 500 µL of PBS, followed by centrifugation at 13’300 rpm for 2.5 minutes. The supernatant was aspirated and 300 µL of undiluted Fixation/Permeabilization Concentrate (Invitrogen) was added into each tube. Pellets were thoroughly resuspended, and cells were fixed at room temperature for at least 20 minutes, prior to being transferred into a new tube and exported from biosafety level 3 conditions. The Fixation/Permeabilization concentrate was diluted by adding 1 mL of eBioscience Fixation/Permeabilization Diluent (Invitrogen) and cells were pelleted at 380 g for 5 minutes at 4°C. Cells were subsequently resuspended in 1 mL of BD CellWASH and stored at 4°C until further analysis. For cell staining, cells were pelleted at 380 g for 5 minutes at 4°C. Primary antibodies (Table 1A) were resuspended in Permeabilization Buffer (Invitrogen), diluted following manufacturer’s instructions. Following centrifugation, the supernatant was aspirated, and the pellets were resuspended in 150 µL of primary antibody solution per tube. Primary antibody staining was allowed to proceed for a minimum of 15 minutes on ice prior to dilution with 1 mL of Permeabilization Buffer followed by centrifugation at 380 g, 4°C for 5 minutes. Secondary antibodies (Table 1B) were diluted in Permeabilization Buffer. After centrifugation, supernatants were aspirated, and cells were resuspended in 150 µL of secondary antibody suspension per tube. Cells were incubated with secondary antibodies for 15 minutes, on ice. Antibody solutions were diluted by the addition of 1 mL of Permeabilization Buffer and cells were pelleted once more at 380 g, 4°C for 5 minutes. Supernatants were aspirated and cells resuspended in 100 µL of BD CellWASH per tube. All samples were filtered using 35 µm cell strainers prior to analysis with a Cytek Aurora flow cytometer equipped with SpectroFlo software. Flow cytometry data was analyzed using FlowJo 10.9.0.

### Production and maintenance of rat brain slice cultures

The preparation of cerebellar OTCs was done as previously reported (*28, 81*). Animal experiments were approved by the Animal Care and Experimentation Committee of the Canton of Bern, Switzerland (licenses BE 6/20 and BE 17/23). Organotypic brain slice culture medium (OTCs medium) was prepared by adding 1% of GlutaMAX and 1% of Antibiotic-Antimycotic to Neurobasal medium (Thermofisher). The dissection medium was produced by adding 6 mg/mL of glucose and 1% of Antibiotic-Antimycotic to Hanks’ Balanced Salt Solution (HBSS) (Thermofisher). Ten-day-old Wistar rats (either from the Central Animal facility of the University of Bern, Switzerland, or Charles River, Germany) were euthanized with an intraperitoneal injection of pentobarbital (Escornarkon, Streuli AG,150 mg/kg body weight). For tissue collection following euthanasia, the head was sterilized with 70% ethanol, the animals were decapitated, the brain was harvested and immersed in ice-cold dissection medium. The cerebellum was isolated from the rest of the brain with a scalpel and bisected along the sagittal plane. The two cerebellar hemispheres were glued onto the cutting plate of a vibratome (Leica VT 1000 S) and cut in parallel with a speed set to 0.40 mm/s, frequency to 10 Hz, and the slice thickness set to 325 μm. The slices were transferred onto the membrane of Transwell inserts (#3450, Sigma Aldrich) (3 slices from different animals per insert) kept on top of ice-cold dissection medium. After all slices were cut, the inserts were transferred to a 6-well plate containing pre-equilibrated OTCs medium freshly supplemented with B27 supplement (Thermofisher) and the plates were incubated at 37°C with 5% CO_2_. The OTCs medium supplemented with B27 was replaced every 2 days.

### Infection of rat brain slice cultures

After 5 days in culture, OTCs were infected. For this purpose, inoculum was prepared by adding viral stock (final concentration 10^5^ TCID_50_/mL) and the respective antiviral at concentrations ranging from 0 to 100 µM to the OTC medium in a final volume of 1 mL per well. Culture medium was removed from the wells and 800 µL of inoculum was added under the inserts, while the remaining 200 μL were evenly distributed on top of the slices in a dropwise manner. The plates were incubated at 37°C with 5% CO_2_ for 1 hour. Following the incubation period, the inoculum was removed from both sides of the inserts. The wells were washed once with 1 mL PBS. One mL of OTCs medium supplemented with B27 (20 μL/mL) and the respective antiviral was added to the bottom of each Transwell insert. Subsequently, the plates were incubated for a period of 3 days, without further medium changes. After 3 days, the total medium was removed and stored at -80°C, and the slices were fixed directly on the inserts with 4% PFA for 90 minutes. Finally, the slices were transferred to PBS and stored at 4°C until further use.

### Immunofluorescence staining of rat brain slice cultures

Fixed samples were initially incubated in blocking buffer (10% FBS, 0.1% Triton X-100 in PBS) for one hour at room temperature. Subsequently, OTCs were incubated with primary antibodies (Table 1A) in blocking buffer for 48 hours on a shaker at 4°C. Following this, three washes for 20 minutes each with PBS were performed, followed by 12-hour incubation in PBS at 4°C. Prior to the subsequent step, a 1-hour incubation in blocking buffer was conducted. Secondary antibodies (Table 1B) in blocking buffer were added, and the samples were incubated for another 24 hours on a shaker at 4°C. At the end of the incubation, the samples were washed with at least 5 changes of PBS over 4 hours at 4°C. Thereafter, the samples were incubated in PBS overnight at 4°C and, ultimately, embedded on a slide using Fluoroshield histology mounting medium (Sigma-Aldrich). Pictures were taken using a fluorescence microscope (Zeiss Axio Imager M1). Brightness and contrast were adjusted equally between experiments using ImageJ 1.53c.

### Use of large language models

During manuscript preparation, the authors used generative AI tools (ChatGPT-5, Nature Research Assistant) to improve readability and clarity, including language editing and grammatical refinement. All content was subsequently reviewed and revised by the authors, who take full responsibility for the final published version.

### Ethics statement

In accordance with Articles 13 and 14 of the Federal Act on Research on Embryonic Stem Cells, and Article 20 of the Ordinance on Research on Embryonic Stem Cells, the work with human H1 ESC line was approved by the Cantonal Ethics Committee of Bern, Switzerland under the authorization number Req-2024-01592 / R-FP-S-2-0023-0000.

## Acknowledgements

This study was supported by the Ministry of Education, Youth and Sports (MŠMT) of the Czech Republic, grant LUAUS25011 within the InterExcellence II program (to L.E) and the Ministry of Health of the Czech Republic in cooperation with the Czech Health Research Council under project No. NU22-05-00659, as well as by the Multidisciplinary Center for Infectious Diseases of the University of Bern.

---

**Supplementary Figure 1:**
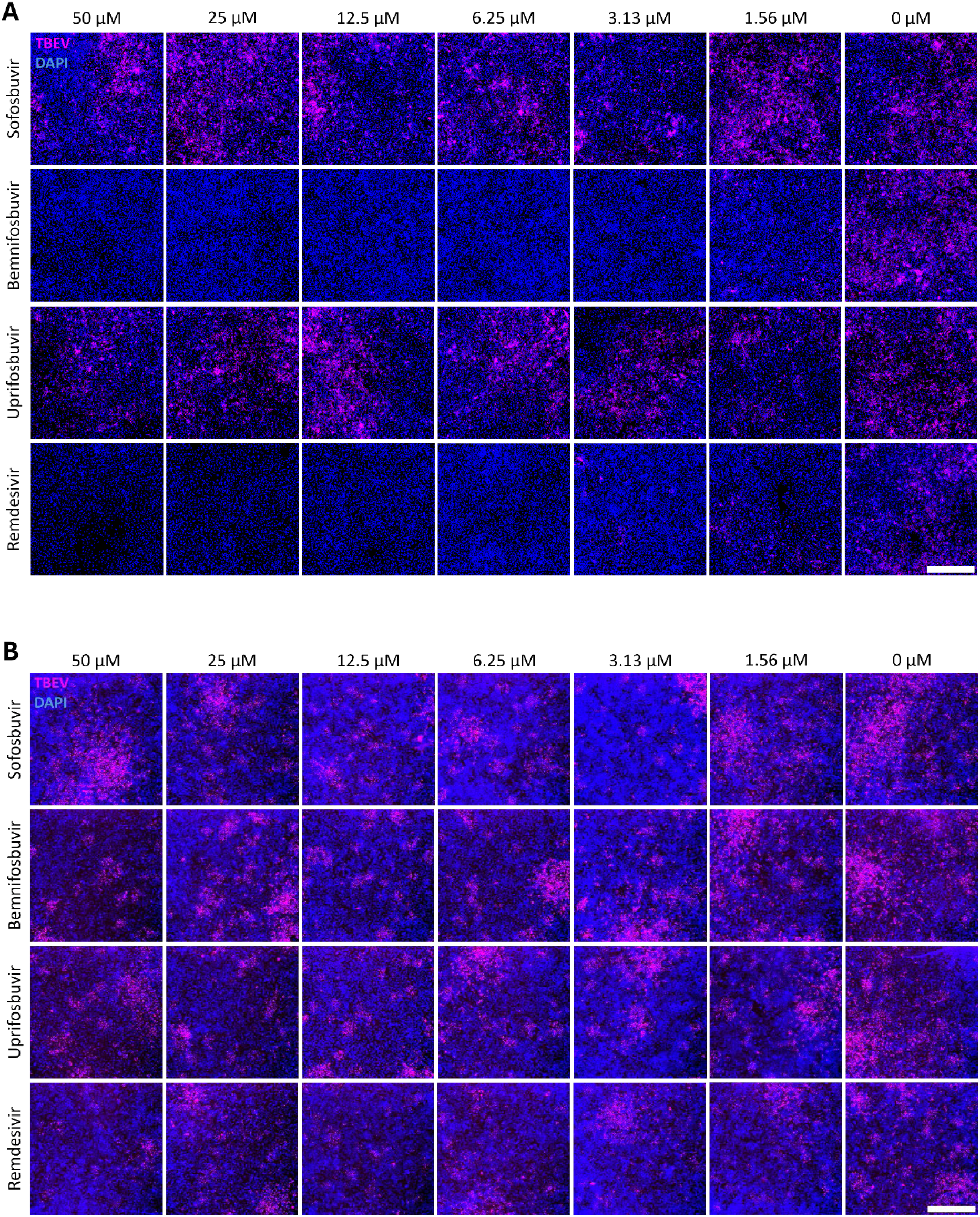
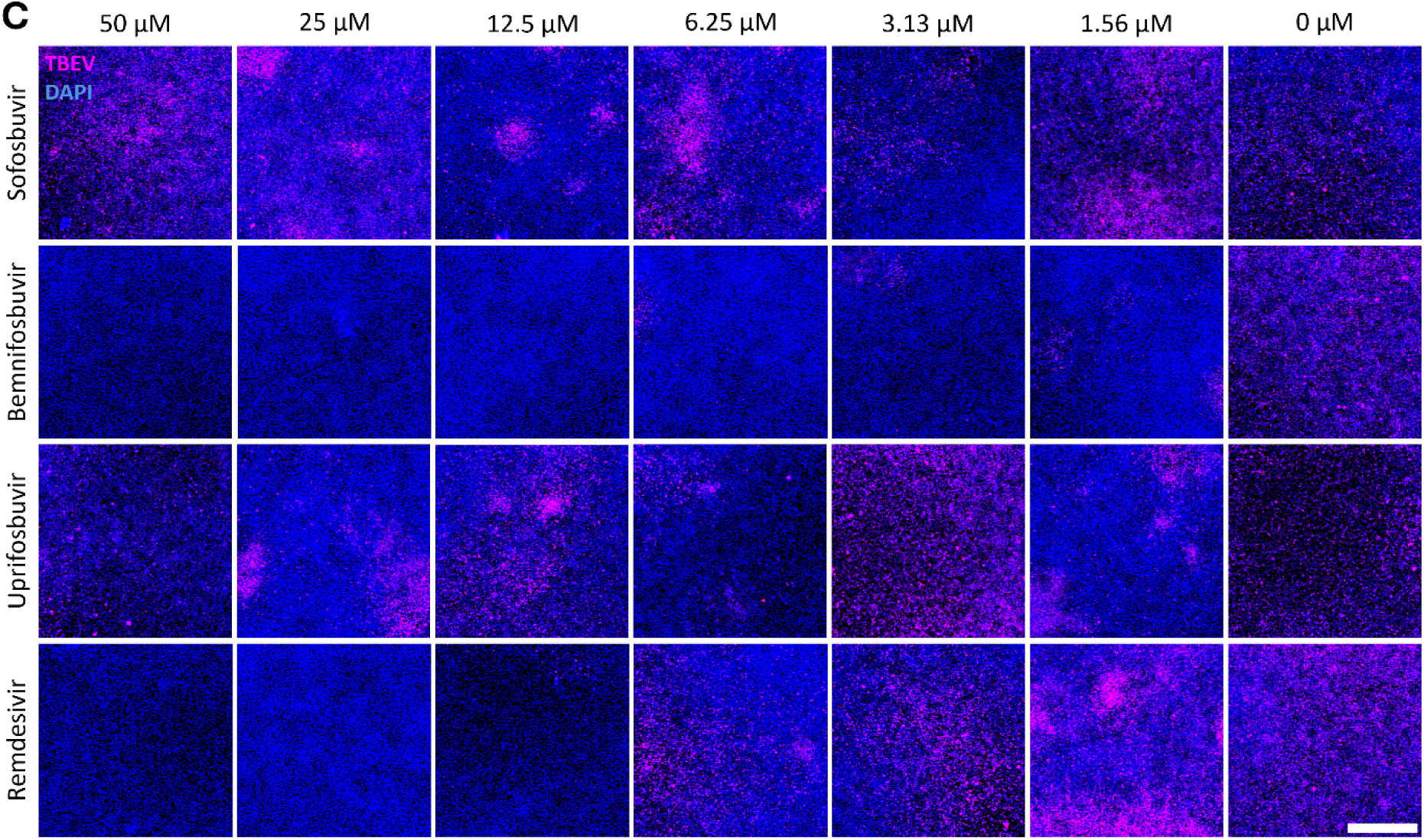
Immunofluorescence staining to monitor TBEV E protein expression in virus-infected, compound-treated cell cultures. A549 (A), Vero (B), and SK-N-SH cells (C) were infected with TBEV and treated with the indicated compounds. At 48 h post infection, the cells were fixed on slides, stained with a mouse orthoflavivirus E-protein–specific antibody (magenta), and counterstained with DAPI (blue). Scale bar, 500 μm.

**Supplementary Figure 2:**
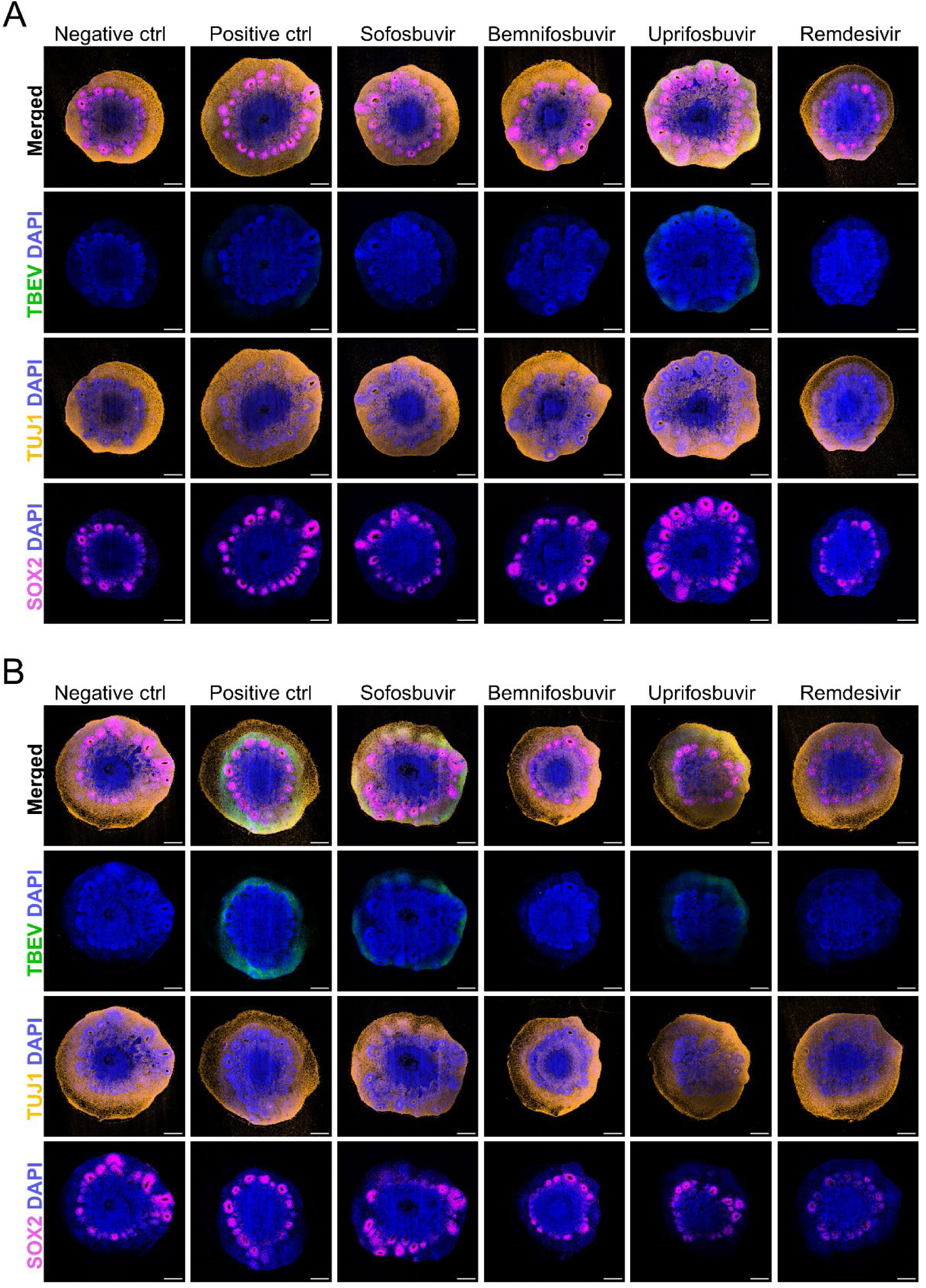
Remdesivir and bemnifosbuvir inhibit TBEV replication at 50 µM. Representative tile-scan confocal micrographs of mock- and TBEV-challenged organoids at 2 days p.i. (**A**) and 4 days p.i. (**B**) with or without treatment with 50 μM of selected NAs. n = 3 independent organoid batches. DAPI, blue; TBEV E-protein (4G2), green; TUJ1, orange; SOX2, magenta. Scale bars, 500 µm.

**Supplementary Figure 3:**
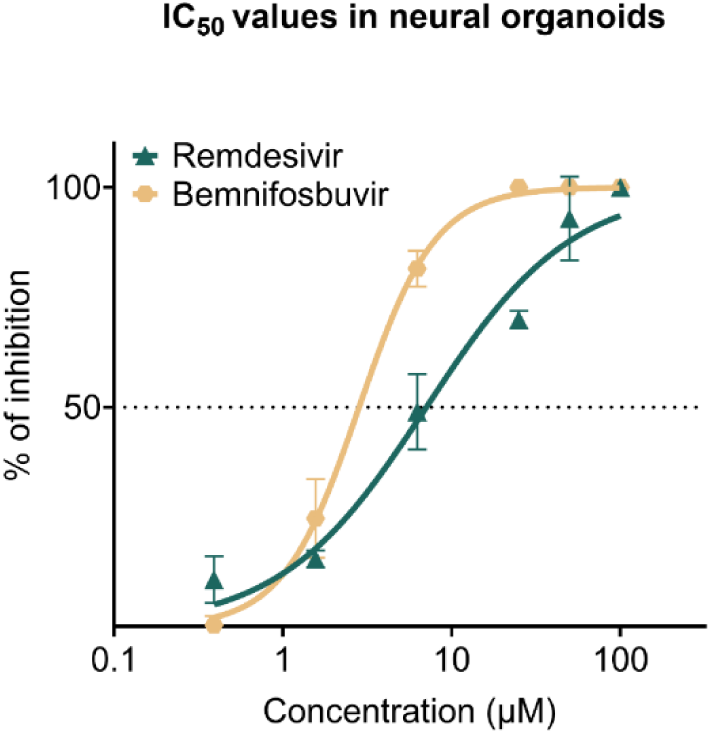
Remdesivir and bemnifosbuvir IC_50_ values in human neural organoids. Mean dose-dependent inhibition of infectious TBEV titers following treatment with remdesivir and bemnifosbuvir at concentrations between 100 μM and 0.39 μM. Dose response curves were obtained using a nonlinear regression method. Points indicate mean values ± s.d. n = 3 independent organoid batches. Horizontal dotted line indicates 50% inhibition.

**Supplementary Figure 4:**
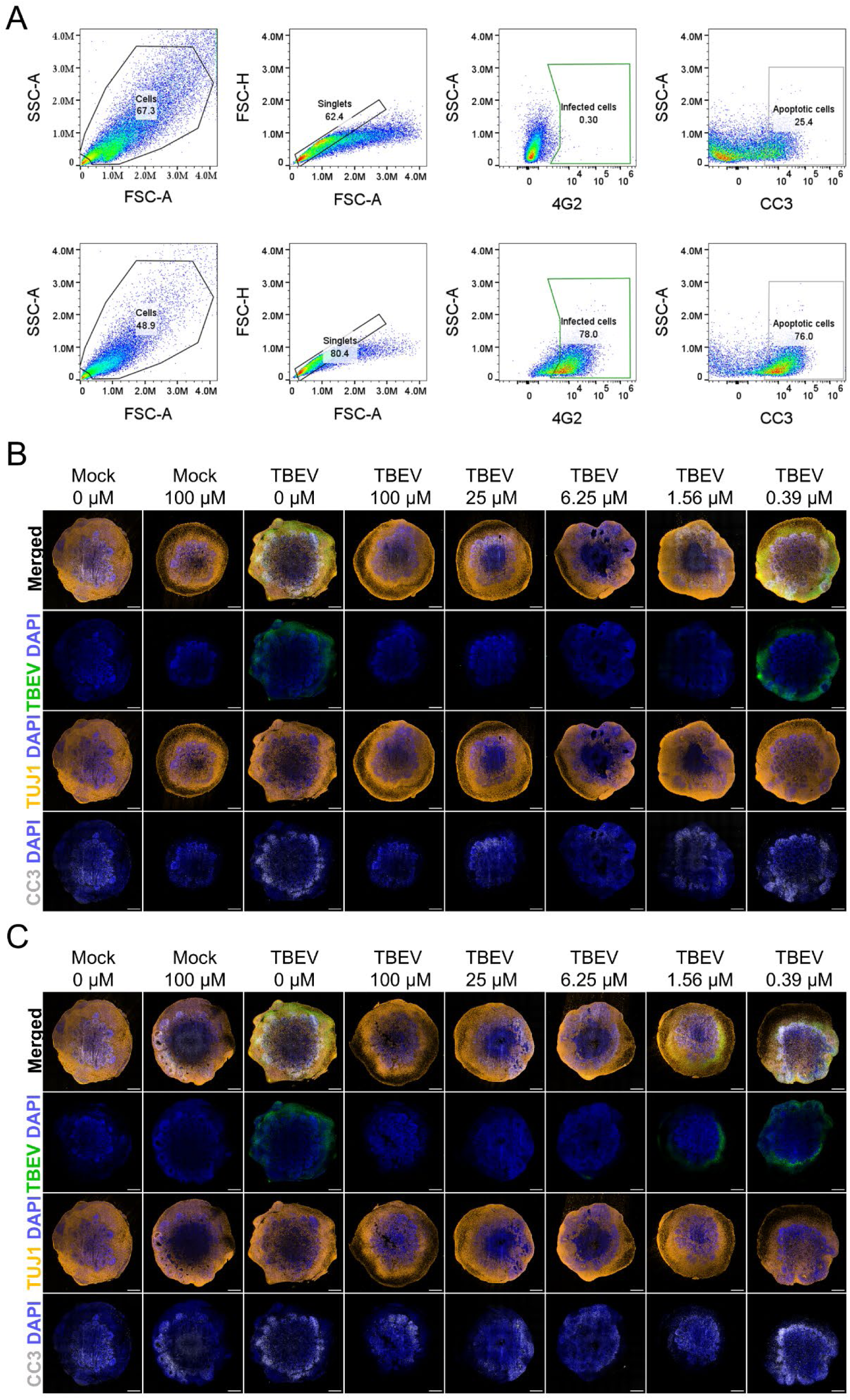
Remdesivir causes increased cell apoptosis in human neural organoids. **A**, Exemplary gating strategy applied to flow cytometry data. The top graphs display a mock-infected and untreated, bottom gates a TBEV-infected and untreated organoid. **B-C**, Representative tile-scan confocal micrographs of mock- and TBEV-challenged organoids with or without treatment with 100 μM-0.39 μM of bemnifosbuvir (**B**) or remdesivir (**C**). n = 3 independent organoid batches. DAPI, blue; TBEV E-protein (4G2), green; TUJ1, orange; CC3, grey. Scale bars, 500 µm.

**Supplementary Table 1:** List of antibodies.

| Primary antibody | Dilution | Host | Catalog # | Source |
| --- | --- | --- | --- | --- |
| TCID<sub>50</sub> assay |  |  |  |  |
| 4G2, clone D1-4G2-4-15 | 1:10 | Mouse | HB-112 | ATCC |
| Immunofluorescence microscopy |  |  |  |  |
| 4G2, clone D1-4G2-4-15 | 1:2 | Mouse | HB-112 | ATCC |
| Calbindin | 1:1000 | Rabbit | AB229915 | Abcam |
| CC3 | 1:500 | Rabbit | 9661 | Cell Signaling Technology |
| T036 | 1:1000 | Human | - | Istituto di ricerca biomedica (Bellinzona) |
| Mouse pan-flavivirus anti-E protein | 1:250 | Mouse | MAB10216-I | Sigma-Aldrich |
| TUJ1 | 1:400 | Chicken | AB41489 | Abcam |
| SOX2 | 1:200 | Rat | 14-9811-82 | Invitrogen |
| Flow cytometry |  |  |  |  |
| 4G2, clone D1-4G2-4-15 | 1:10 | Mouse | HB-112 | ATCC |
| CC3 | 1:800 | Rabbit | 9661 | Cell Signaling Technology |

| Secondary antibody | Dilution | Host | Catalog # | Source |
| --- | --- | --- | --- | --- |
| TCID<sub>50</sub> assay |  |  |  |  |
| Rabbit anti-mouse HRP | 1:250 | Rabbit | P0260 | Dako |
| Immunofluorescence microscopy |  |  |  |  |
| Cy3 AffiniPure Donkey Anti-Human | 1:1000 | Donkey | 709-165-149 | Jackson ImmunoResearch |
| Donkey anti-Rabbit IgG, Alexa Fluor 488 | 1:1000 | Donkey | A21206 | ThermoFisher |
| Goat anti-Chicken IgY, Alexa Fluor 546 | 1:200 | Goat | A-11040 | Invitrogen |
| Goat anti-Mouse, Alexa Fluor 647 | 1:1000 | Goat | A-21235 | Invitrogen |
| Goat anti-Mouse IgG2a, Alexa Fluor 488 | 1:200 | Goat | A-21131 | Invitrogen |
| Goat anti-Rabbit IgG, Alexa Fluor 647 | 1:200 | Goat | A-21244 | Invitrogen |
| Goat anti-Rat IgG, Alexa Fluor 647 | 1:500 | Goat | A-21247 | Invitrogen |
| Flow cytometry |  |  |  |  |
| Goat anti-Mouse IgG2a, Alexa Fluor 488 | 1:500 | Goat | A-21131 | Invitrogen |
| Goat anti-Rabbit IgG, Alexa Fluor 647 | 1:500 | Goat | A-21244 | Invitrogen |
| SSEA-5 | 1:50 | Mouse | 60063AZ | Stem Cell Technologies |
| TRA-1-60 | 1:50 | Mouse | 60064PE | Stem Cell Technologies |

List of primary and secondary antibodies used for TCID<sub>50</sub> assay, immunofluorescence microscopy and flow cytometry analysis.

## Notes

### Competing Interest Statement

The authors have declared no competing interest.

